# Universal chimeric Fcγ receptor T cells with appropriate affinity for IgG1 antibody exhibit optimal antitumor efficacy

**DOI:** 10.1101/2022.10.23.513394

**Authors:** Wen Zhu, Yang Wang, Liangyin Lv, Hui Wang, Wenqiang Shi, Zexin Liu, Mingzhe Zhou, Jianwei Zhu, Huili Lu

## Abstract

Developing universal CARs with improved flexible targeting and controllable activities is urgently needed. While several studies have suggested the potential of CD16a in tandem with monoclonal antibodies to construct universal CAR T cells, the weak affinity between them is one of the limiting factors for efficacy. Herein, we systematically investigated the impact of Fcγ receptor (FcγR) affinity on CAR T cells properties by constructing universal CARs using Fcγ receptors with different affinities for IgG1 antibodies, namely CD16a, CD32a, and CD64. We demonstrated that the activities of these universal CAR T cells on tumor cells could be redirected and regulated by IgG1 antibodies. In xenografted mice, 64CAR chimeric Jurkat cells with the highest affinity showed significant antitumor effects in combination with herceptin in the Her2 low expression U251 MG model. However, in the CD20 high expression Raji model, 64CAR caused excessive activation of CAR-T cells, which resulted in cytokine release syndrome (CRS) and the decline of antitumor activity, and 32CAR with a moderate affinity brought the best efficacy. Our work extended the knowledge about FcγR-based universal CAR T cells and suggested that only the FcγRCAR with an appropriate affinity can offer the optimal antitumor advantages of CAR T cells.

**Graphical abstract:** 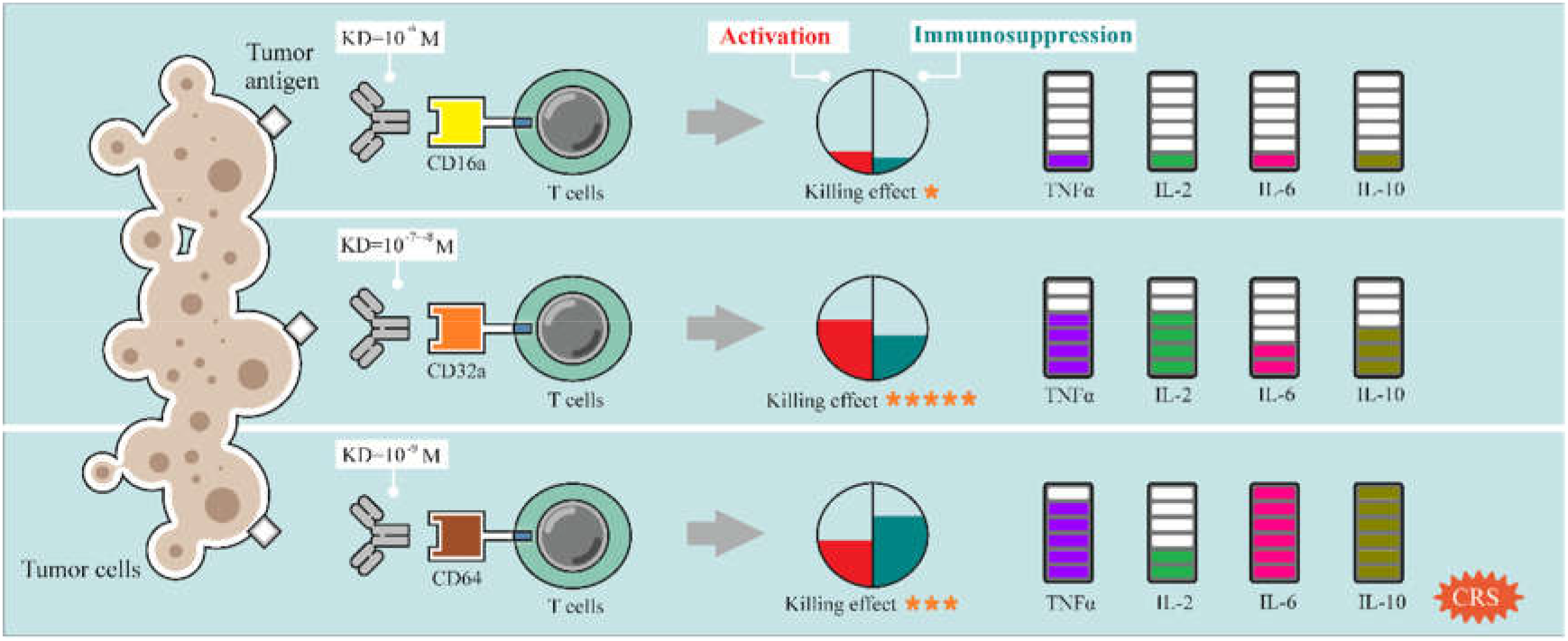

Universal CAR T cells based on Fcγ receptors exhibit a specific tumor-killing effect. However, the affinities of Fcγ receptors greatly influence the efficacy and adverse effects *in vivo*.

## 1. Introduction

Chimeric antigen receptor (CAR) T cells have shown remarkable effectiveness in treating some hematological malignancies, leading to the recent FDA approvals of several CD19-targeting CAR T-cell therapies^1^. CARs are usually composed of extracellular single-chain fragment variable (scFv) antibody and intercellular signaling domains, which bestow T cells with the capacity to recognize antigens and receive activation and co-stimulatory signals, thus exhibiting potent antitumor activity^2^. Though achieving impressive success in treating hematological malignancies, CAR T cells often fail to mount an effective response in solid tumors, with numerous cases of toxic side effects^3^. In solid tumor patients, both the over-activation of CAR T cells in the tumor site and the recognition of low levels of antigen expressed on the normal cells can cause fatal CRS^4^. The infiltrating CAR T cells are also particularly sensitive to the chronic inflammatory microenvironment of solid tumors, in which the immunosuppressive context can quickly induce dysfunctional states of T cells^5^. Furthermore, clinical-grade CAR T cells are technically challenging and require extensive engineering and safety testing, which hinder the wide clinical application^6^.

The extracellular scFv domain may play an important role in the problems above. Several intrinsic properties of scFv impede the development of CAR T-cell therapy. No single tumor associated antigen (TAA) is expressed by all cancer types, so scFv encoded by CARs need to be constructed for each potential TAA. In addition, the inability of the current scFv-based CARs to target more than one antigen on tumors may lead to the preferential growth of more aggressive, antigen-negative cancer cells. To solve this, universal CARs that bind to an adaptor but do not directly recognize the target antigen were developed^7^. Among these dual-element systems, chimeric anti-tag scFv and tagged adaptors occupy major part, such as “anti-FITC scFv & FITC labeled antibodies”^8^, “anti-peptide neo-epitope (PNE) scFv & PNE labeled antibodies”^9, 10^, and “anti-5B9 tag scFv & 5B9 tag labeled scFv”^11, 12^. The combination of universal CARs with such multi-specific adaptors gives a chance to target a diversity of antigens simultaneously without the need for re-engineering T cells.

However, scFv-based universal CARs face some unmanageable shortages. Due to the sequence heterogeneity, the N terminal distributed scFv makes a profound but enigmatic impact on the expression level of CARs^13^. Moreover, the tendency to form aggregated scFv on the membrane resulted in the self-activation without antigen and subsequent activation-induced cell death (AICD)^14^. What’s worse, the immune responses against murine-derived scFv also caused even death in the clinic^15^. Last but not least, the high affinity of scFv (~nM) may elicit a potent T cell signaling and subsequent significant cytokine response, with unpredictable severity^16^. Targeting TAAs expressed at low-to-intermediate levels on normal tissues with CAR T cells raises safety concerns about on-target off-tumor toxicities^17^. Some researchers achieved promising outcomes by simply utilizing a low-affinity scFv. By decreasing the affinity of ErBb2 targeted 4D5 from KD=0.3 nM to 1119 nM, Liu et al. proved that the low-affinity CARs demonstrated more discrimination between target cells with high or low target expression levels^18^. Esther et al. showed that when a CD28 costimulatory domain and a 41BB intracellular domain were used together, the CD38 CARs of very low affinity (KD<1.9 μM) mediated superior cytotoxicity^19^.

Since the work of optimal affinity screening for each scFv is highly time and labor-consuming, it’s an ideal approach to substitute scFv with an alternative molecule with a simple and stable structure to address the limitations of scFv-based universal CAR T cells. Giuseppe et al. set an Fc gamma chimeric receptors (Fcγ-CRs)-based strategy, where the scFv was replaced with the extracellular domain of FcγRIIIA (CD16a), thus T cells targeting can be mediated by clinically approved monoclonal antibodies (mAbs)20, 21. This scheme has many intriguing features. In combination with mAbs, CD16a-CR T cells exhibit anticancer activity by a widely known antibody-dependent cellular cytotoxicity (ADCC) mechanism. Provided that mAbs with the appropriate specificity to TAAs are available in the market, CD16a-CR T cells can target multiple cancer types. One can shut down the activity of CD16a-CR T cells by simply withdrawing the administration of mAb or adjusting the plasma concentration of mAb. About a dozen studies show that CD16a-CR T or NK cells could inhibit the growth of tumor cells *in vivo* when concomitantly infused with mAbs^22–29^. Several clinical trials (NCT03189836, NCT02776813, and NCT03266692) were also launched to evaluate the combination of CD16a-CR T cells and antibody. However, scattered studies explored other Fcγ receptors, which can bind antibodies with different affinities and possess potentials for CAR T cells mediation.

In the current study, we constructed the universal CAR T cells armed with three Fcγ receptors with different antibody affinities with KD values ranging from of 10^−6^ M to 10^−9^ M, namely CD16a, CD32a, and CD64, and systemically examined their efficacy and safety profiles to explore the influence of CAR affinity on the antitumor efficacy of CAR T cells. While positive correlations between effector cells’ activities and affinities were observed *in vitro*, the *in vivo* experiments demonstrate an intricate influence of affinity. 16CAR failed to inhibit tumor growth for the weak affinity. Although exhibiting the best tumor inhibition effect in the U251 MG model with low Her2 density, 64CAR strong affinity caused CRS and the following immunosuppression state of CAR T cells in the Raji model with high CD20 density, where 32CAR with a moderate affinity displayed considerable antitumor activity. In summary, this work firstly reported the side-by-side comparison of the three Fcγ receptors in FcγRCAR T cells, expanding this promising universal CAR T therapy’s clinical versatility and suggesting the pivotal role of affinity.

## 2 Material and Methods

### 2.1 Cells

All the cell lines in this study were purchased from American Type Culture Collection (ATCC). The HEK293T cell line used for the production of infectious lentivirus was maintained in DMEM supplemented with 10% fetal bovine serum (FBS) (Gibco, USA). The HEK293E cell line used for the production of rituximab and its variants was maintained in SMM 293-TII Expression Medium (SinoBiological, China). The human immortalized T cell line Jurkat, CD20^+^ lymphoma cell line Raji, Her2^+^ human breast cancer cell line BT474, and Her2^+^ human glioblastoma cell line U251 MG were cultured in RPMI-1640 supplemented with 10% FBS. Cells were cultured at 37 °C under 5% CO_2_ and 95% humidity. The cell lines were passaged for three to eight times before use, or kept in culture for a maximum of 3 weeks.

### 2.2 Plasmids construction, virus production, and gene transduction

The pRRLSIN-pMD2.G-psPAX2 three plasmids lentivirus system were purchased from Addgene. The FCGRIIIA-F (CD16a, UniProt: P08637), FCGRIIA-H (CD32a, UniProt: P12318), and FCGRI (CD64, UniProt: P12314) cDNA were obtained by PCR using the cDNA of human peripheral blood mononuclear cell as template. The EF1α-core promoter (GenBank: MW560964.1), CD8α signal peptide (UniProt: P01732), the hinge, transmembrane, and costimulatory domain of CD28 (UniProt: P10747), the costimulatory domains of CD137 (UniProt: Q07011) and CD247 (UniProt: P20963) were cloned from another plasmid previously made in our laboratory. These constructs were assembled by overlapping extension PCR. The constructs and expression cassette were sub-cloned into the pRRLSIN vector by Gibson Assembly. Genes and amino acid sequences of CARs were supplied in the supplementary file.

For lentivirus production, HEK293T cells were transfected with pMD2.G, psPAX2, and pRRLSIN transfer plasmid using Exfect Transfection Reagent (Vazyme, China). Supernatants containing virus particles were collected approximately 36 and 60 h post-transfection and concentrated with lentivirus concentration reagent (Genomeditech, China). The lentivirus titers were determined by transducing HEK293T cells with serial dilutions.

Jurkat cells were grown to the mid-log phase before transduction. For each CAR, 2E5 cells were given lentivirus (multiplicity of infection = 10) in RPMI-1640 with 10% FBS and Hitrans G pro-infection kit (Genechem, China) to a final volume of 200 μL, and incubated in a 48-well plate for 12 h. After further 72 h culture, the cells were analyzed. Photographs were captured with an Olympus CKX53 fluorescence microscope (Olympus, Japan). Western Blot was performed with anti-human CD247 antibody. And cells were stained for flow cytometry with anti-human CD16a, anti-human CD32a, and anti-human CD64 antibodies (APC) using a CytoFlex S (Beckman Coulter, USA). All the antibodies were purchased from SinoBiological, China.

### 2.3 Antibody binding and cell aggregation assays

To measure the FcγRCARs’ antibody-binding capacity, we incubated 2E5 JK-EGCAR/FcγRCAR cells with different concentrations of herceptin (Hct) (Henlius, China) in 200 μL PBS at 4 °C for 30 min. After washing twice with PBS, cells were incubated with goat anti-human immunoglobulin G (IgG) antibody (APC) (Yeasen, China) at 4 °C for 20 min, and cell staining was measured by fluorescence activated cell sorting (FACS). To further verify the binding capacities of these JK-FcγRCAR cells to other IgG1 antibodies, the experiments were repeated with human IgG under the same conditions.

To determine whether antibody can mediate the aggregation of FcγRCAR T cells and tumor cells, we labeled CD20-positive Raji cells with DiI (red) (Beyotime, China), and then incubated them with excessive rituximab (Rtx) (Henlius, China) at 4 °C for 30 min. JK-EGCAR/FcγRCAR cells were labeled with DiO (green) (Beyotime, China). After washing twice with PBS, 2E4 Raji cells and 2E5 JK-EGCAR/FcγRCAR cells were mixed to a final volume of 200 μL PBS and transferred to a 2 mL EP tube. The tubes were shaken softly on a transference de-coloring shaker at room temperature for 60 min. Then the cells were examined for aggregation and photographed by fluorescence microscope, and the proportions of aggregated Raji cells (PE-FITC double-positive/PE positive) were determined by FACS.

To investigate the block of human serum IgG on FcγRCARs, we incubated JK-FcγRCAR cells in human serum, PBS with 10% human serum, or PBS for 30 min, respectively. After washing twice with PBS, cells were incubated with goat anti-human IgG antibody (APC), and fluorescence was measured by FACS. To further explore whether such block would interfere the recognition of FcγRCARs to mAb opsonized target cells, we incubated JK-FcγRCAR-DiO cells in human serum for 30 min, then mixed them with Raji-DiI cells pre-incubated with Rtx or not. After 3 h of co-incubation, cell aggregation was detected by FACS and fluorescence microscope, respectively.

### 2.4 ADCC, cell activation, cytokine release, and cell proliferation assay

The ADCC activities mediated by the JK-EGCAR/FcγRCAR cells were detected by standard L-lactate dehydrogenase (LDH)-release assay. Briefly, 1E4 Raji cells were incubated with 5E4 JK-EGCAR/FcγRCAR cells at various concentrations of Rtx in 100 μL RPMI-1640 in 96-well plats (Biofil, China). After 4 h of incubation, the supernatants were collected, and the concentrations of LDH were measured with LDH Cytotoxicity Assay Kit (Beyotime, China). The percentage of specific lysis was calculated as: (experimental release - spontaneous release) / (maximal release -spontaneous release) * 100 (%).

Cell activation, cytokine release, and cell proliferation were measured under a saturated concentration of Rtx. 5E4 JK-EGCAR/FcγRCAR cells, 1E4 Raji cells, and 0.1 μg/mL Rtx were incubated in 100 μL RPMI-1640 for 4 h. Then the cells were harvested and analyzed with mouse anti-human CD69/CD107a antibody (APC) (SinoBiological, China) by FACS. In other experiments, FBS was added to a final concentration of 10%, and the incubation time was elongated to 48 h, then the levels of TNFα, IFNγ, and IL-2 in culture supernatants were measured using enzyme-linked immunosorbent assay (ELISA) kits (Multi Sciences, China). For cell proliferation assay, 5E4 carboxyfluorescein succinimidyl ester (CFSE) stained JK-EGCAR/FcγRCAR cells, 1E4 Raji cells, and 0.1 μg/mL Rtx were incubated in 100 μL RPMI-1640 with 10% FBS for 24 h, and then the JK-EGCAR/FcγRCAR cells were harvested and analyzed by FACS.

All these experiments were also performed the same with BT474 cells and Hct as the target cells and adaptor, except that the cell number of BT474 cells and JK-EGCAR/FcγRCAR cells were 3E3 and 15E3, respectively.

### 2.5 De-glycosylation of antibody

To remove the attached sugars, 10 μg Rtx and Hct were digested by 500 unit PNGase F (NEB, USA) at 37 °C for 24 h under native conditions. The digestion products were analyzed by reducing SDS-PAGE. Antibody binding, cell aggregation, and ADCC assay were performed exactly the same as the previous section, except that the primary Rtx and Hct were replaced by the de-glycosylated ones.

### 2.6 In vivo antitumor activity mediated by intravenously infused JK-FcγRCAR cells and antibody in a xenografted mouse mode

All *in vivo* experiments were conducted following guidelines from the Institutional Animal Care and Use Committee of Shanghai Jiao Tong University. Six-week-old BALB/c-nu (nude) female mice (Shanghai Slac Laboratory Animal) were subcutaneously inoculated with 5E6 U251 MG or 4E6 Raji cells in 50 μL PBS mixed with 50 μL Matrigel (Corning, USA), and the day was recorded as day0. From day5, the tumor volumes were detected every day with a vernier caliper and calculated as length*width*width/2, and the body weights and physical status were also recorded. When the tumor reached volumes of around 200 mm^3^ (day8 for the U251 MG model and day10 for the Raji model, respectively), 3E6 JK-EGCAR/FcγRCAR cells (in 100 μL PBS) or 100 μL PBS were intravenously administered. Then 20 μg Rtx/Hct (in 100 μL PBS) or 20 μg human IgG (in 100 μL PBS) were intravenously administered every other day (day9/11/13/15 for the U251 MG model and day11/13/15/17 for the Raji model, respectively). At the end point (day16 for the U251 MG model and day18 for the Raji model, respectively), after about 500 μL of blood was collected from the venous sinus to anticoagulation tubes, the mice were sacrificed, and the spleens and tumors were also collected.

The lymphocytes of blood and spleens were isolated with Mouse 1× Lymphocyte Separation Medium (Dakewe, China). For tumor lysis, 100 mg tumor tissue was digested with 1 mg/mL collagenase IV and 0.6 mg/mL hyaluronidase. The tubes were shaken in a shaker at 150 rpm and 37 °C for 60 min. The lysis solutions were filtered with a 200 mesh nylon net, and the flow-through was centrifuged at 300 g for 5 min. After the supernatants were discarded, the cells were re-suspended with 6 mL PBS and filtered with a 200 mesh nylon net. If necessary, Red Blood Cell Lysis Buffer (Beyotime, China) was applied to remove erythrocytes. The single-cell suspensions of blood, spleens, and tumor tissues were incubated with TruStain FcX™ (anti-mouse CD16/32 antibody) (Biolegend, USA) and Human TruStain FcX (Biolegend, USA), subsequently anti-mouse CD45.2 antibody (PE) (Biolegend, USA), anti-human CD3e antibody (APC) (SinoBiological, China). For the single-cell suspensions of tumor tissues, anti-human CD25 antibody (PE) (SinoBiological, China) or anti-human PD-1 antibody (PE) (SinoBiological, China) were also added. After washing twice with PBS, the cells were analyzed by FACS.

Immediately after collection, 100 μL of blood were transferred to new tubes, and the plasma was obtained by centrifugation at 1000 g for 10 min using a refrigerated centrifuge. Tumor and serum samples were stored at −80 °C until further analysis. Tumors (100 mg) were macerated in liquid nitrogen, dissolved in 500 μL PBS and centrifuged at 3000 rpm for 10 min. Supernatants were transferred to new tubes, and the concentrations of cytokines in tumor and serum were determined with ELISA kits (Multi Sciences, China).

### 2.7 Expression, purification, and affinity detection of the rituximab variants

The expression plasmids of Rtx variants were constructed according to the previous report^30^ and prepared using the Endo-Free Plasmid Maxi Kit (Omega Bio-Tek, Norcross, GA, USA). HEK293E cells were seeded at 5E5 cells/mL in SMM 293-TII Expression Medium and adjusted to 1E6 cells/mL 24 h later. The plasmids (0.5 μg/1E6 cells) were diluted with 150 mM NaCl to a concentration of 40 μg/mL and mixed with 40-kDa polyethyleneimine (PEI, Polysciences, Warrington, PA, USA) (DNA: PEI = 1:4, w/w). After incubation for 15 min at room temperature, the mixture was added to the cell culture. The culture supernatants were harvested 6 days post-transfection, and the antibodies were purified by protein A affinity chromatography (GE Healthcare Life Sciences) as previously described^31^. All purified proteins were ultra-filtrated in PBS, sterilized by filtration using a 0.22 μm filter, and frozen in −80 °C freezer until use.

To detect the affinities of Rtx variants to CD20, the ELISA plate was coated with 0.5 μg/mL human CD20 at 4 °C overnight and blocked with 1% (w/w) BSA for 2 h at room temperature, then various concentrations of Rtx-L/M/H and commercial rituximab were added and allow to bind for 2 h at room temperature. After washing, goat anti-human IgG antibody (HRP) (Beyotime, China) was added, and the plate was incubated for 1 h at room temperature. After washing, tetramethylbenzidine (TMB) (Solarbio, China) was added, and the reaction was finished with H_2_SO_4_ (2 M). The OD450 values were detected using a Synergy LX multi-mode reader (BioTek, USA), and the data were fitted using sigmoidal fitting equation in Origin Pro 8.5 software.

### 2.8 Statistical analysis

Data are shown as mean ± standard deviation (SD), and statistical analysis was based on two-tailed heteroscedastic Student’s t-test. Statistical significance is considered at p<0.05. When differences are statistically significant, the significance is represented with asterisks (*) according to the following values: p<0.05 (*), p<0.01 (**), and p<0.001 (***).

## 3. Results

### 3.1 Construction and detection of the chimeric Fcγ receptors

CAR constructs composed of Fcγ receptor as the outer-membrane domain were generated in this study (Fig. 1A). To test the influence of affinity on the therapeutic properties of FcγRCAR T cells, the outer-membrane domains of FCGRIIIA-F/FCGRIIA-H/FCGRI (CD16a-F/CD32a-H/CD64) genes were cloned and fused with the hinge, transmembrane, and costimulatory domain of CD28, the costimulatory molecule CD137 (4-1BB), and the stimulatory molecule CD247 (CD3ζ), in tandem with P2A peptide-linked EGFP (Fig. 1B). Another construct in which EGFP replaced the Fcγ receptor was constructed as the negative control. For the proof of concept, Jurkat cell lines were used as the effector cells and transduced with the pRRSLIN-EGCAR/16CAR/32CAR/64CAR lentiviruses. Approximately 72 h post-transduction, the membrane chimeric and cytoplasm distributed EGFP in JK-EGCAR, and JK-FcγRCAR cells respectively were observed with a fluorescence microscope (Fig. S1). The CAR proteins were also detected by Western Blot and FACS (Fig. 1C and 1D). The expression levels of the three FcγRCARs on Jurkat cells were comparable, whereas the EGCAR was higher. The larger molecular weights of the three FcγRCARs than the theoretical ones infer different glycosylation modes. The positive ratios of the three FcγRCARs were high enough (91.1%, 74.3%, and 95.7%) for the subsequent experiments. Though the lentivirus infection process bought an irreversible adverse effect on the cells, there was no notable difference in proliferation activity between the four groups (Fig. S2).

**Figure 1.**
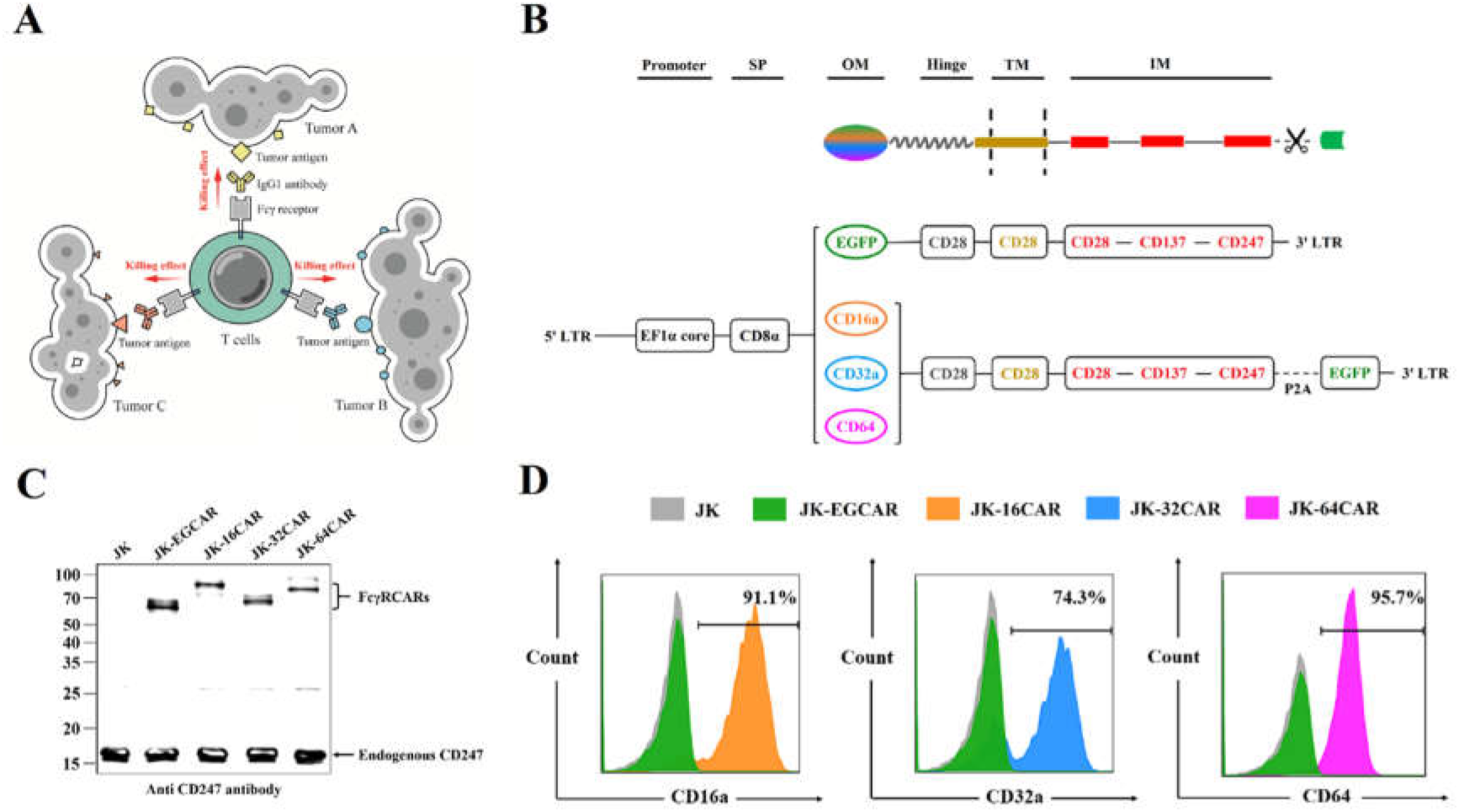
Construction and detection of the chimeric Fcγ receptors. (**A**) Illustration of the universal CAR T therapy based on Fcγ receptors and antibodies. (**B**) Construction of the transfer plasmids of EG/16/32/64CAR. (**C**) Western blot analysis of the EG/16/32/64CAR in Jurkat cells with anti-human CD247 antibody. Lysates of 1E5 cells were loaded per well. (**D**) FACS analysis of the FcγRCARs on Jurkat cells’ membrane with anti-human CD16a, CD32a, and CD64 antibody (APC).

### 3.2 Antibody-binding capacity of different FcγRCARs

The antibody-binding capacities of the FcγRCARs were explored by incubating the JK-EGCAR/FcγRCAR cells with different concentrations of Hct and excessive (10 μg/mL) goat anti-human IgG antibody (APC). The final APC fluorescence intensity characterized the number of bound Hct, reflecting the affinity of the FcγRCARs to antibody (Fig. 2A). As shown in Fig. 2B, at any given concentration of Hct, Jurkat cells transduced with the receptor with higher affinity had a more vigorous APC fluorescence intensity, indicating a higher antibody-binding capacity. The concentration of Hct at which the three FcγRCARs displayed a noticeable shift on the APC axis was 1, 0.1, and 0.01 μg/mL for 16CAR, 32CAR, and 64CAR, respectively (Fig. 2C). No shift was observed in the EGCAR group even at 10 μg/mL Hct, confirming that the chimeric FcγRCARs mediated antibody-binding properties. The 64CAR displayed no more shift when the concentration of Hct was higher than 0.1 μg/mL, indicating that it is the saturated antibody concentration for the three FcγRCARs. The different affinities could also be speculated from the different baseline shifts in the group of none Hct and 10 μg/mL goat anti-human secondary antibody. We got similar results with human IgG (Fig. 2D and 2E), which demonstrated that the three FcγRs in CAR constructs maintained their intrinsic binding properties to IgG1 antibodies.

**Figure 2.**
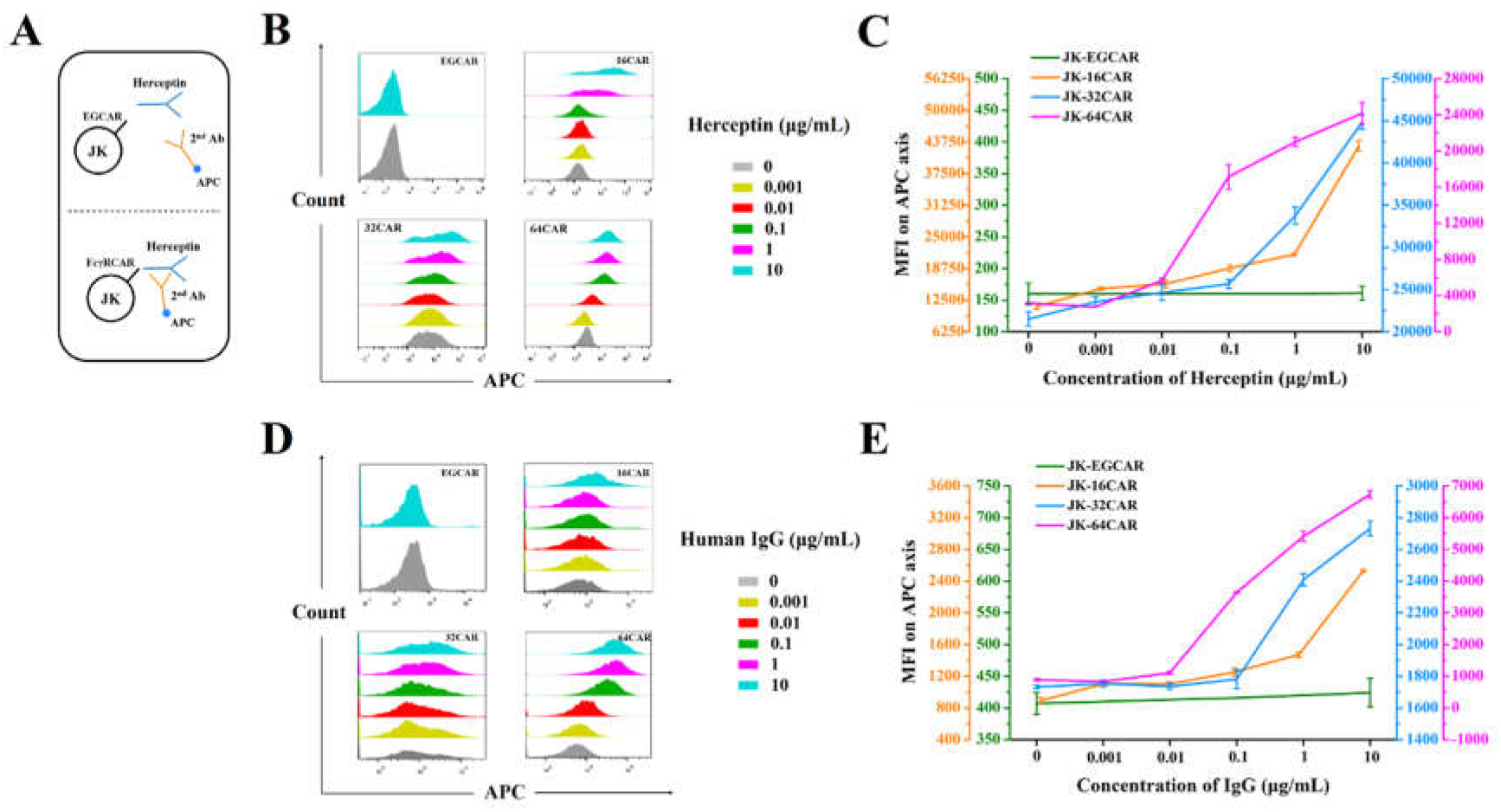
Antibody-binding capacities of the FcγRCARs. (**A**) Scheme of the antibody-binding capacity assay. (**B**) FACS analysis of the bound antibodies on the FcγRCARs. JK-EGCAR/FcγRCAR cells were incubated with different concentrations of herceptin and goat anti-human IgG antibody (APC) in sequence, and then cell staining was measured by FACS. (**C**) MFI of the bound herceptin on JK-EGCAR/FcγRCAR cells. (**D**) FACS analysis of the bound human IgG on the FcγRCARs. JK-EGCAR/FcγRCAR cells were incubated with different concentrations of human IgG and goat anti-human IgG antibody (APC) in sequence, and then cell staining was measured by FACS. (**E**) MFI of the bound human IgG on JK-EGCAR/FcγRCAR cells.

### 3.3 Binding of JK-FcγRCAR cells to antibody opsonized tumor cells

Cell aggregation assays were performed to verify the ability of the JK-FcγRCAR cells to bind antibody-opsonized tumor cells. Raji (CD20^+^) and Rtx were used as the model tumor cell and antibody, respectively. The JK-EGCAR and JK-FcγRCARs cells were stained with fluorescent dye DiO to cover the weak, uneven, and unstable expression of EGFP (15-50% positive ratio, data not shown), then mixed with Raji-DiI or Raji-DiI-Rtx cells at an E: T ratio of 10 for 60 min, finally observed with a fluorescence microscope and measured the DiI-DiO doublets by FACS (Fig. 3A).

**Figure 3.**
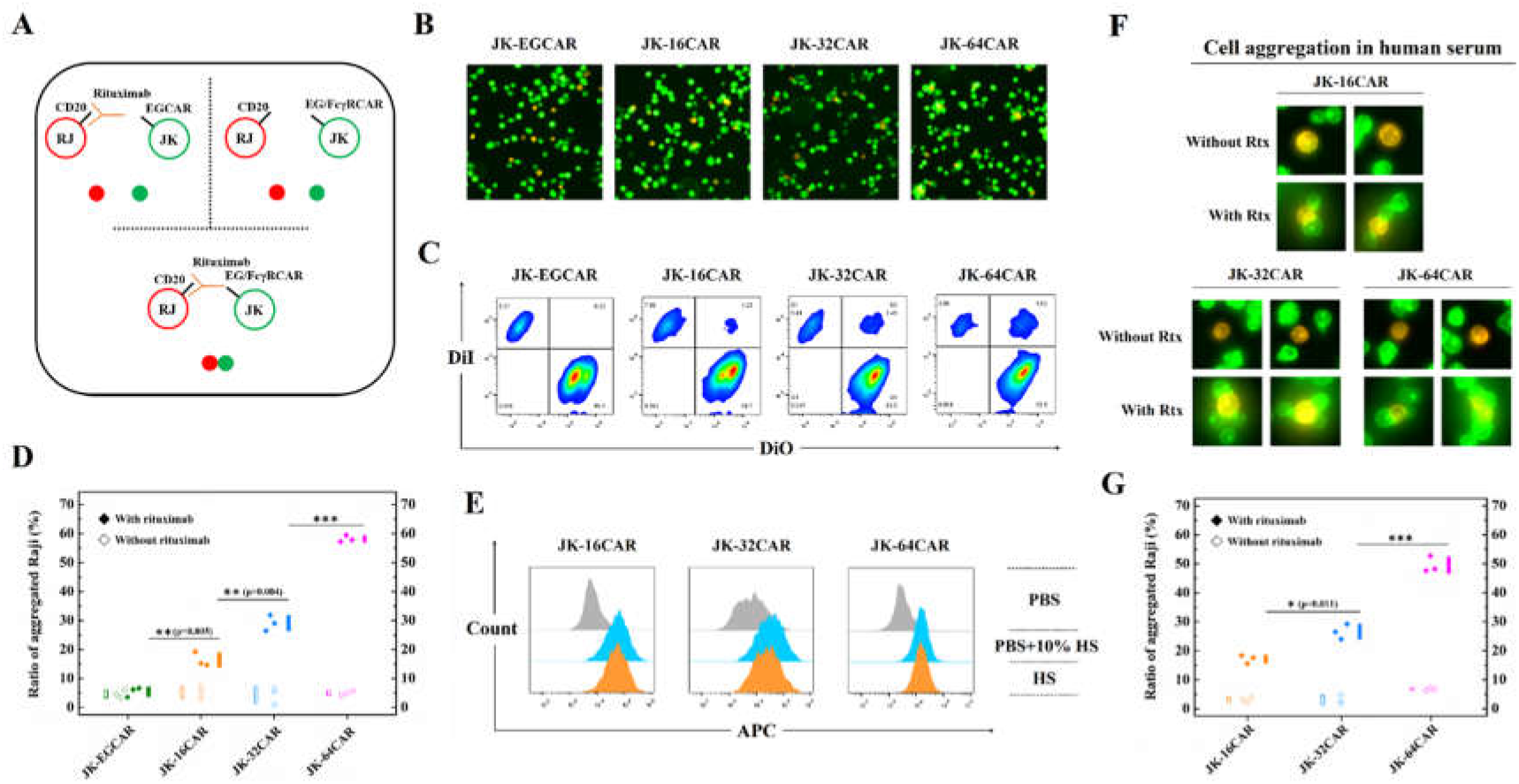
Binding of JK-FcγRCAR cells to antibody opsonized tumor cells. (**A**) Scheme of the antibody opsonized tumor cells-binding capacity assay. JK-EGCAR/FcγRCAR cells were labeled with DiO (green), and Raji cells were labeled with DiI (red). The two cells were mixed (E: T ratio=10) with or without rituximab. (**B**) FACS analysis of the ratio of aggregated Raji cells after 1 h co-incubation in PBS. The aggregated Raji cells were shown as the PE-FITC double-positive events. (**C**) The detection of cell aggregation in PBS with a fluorescence microscope. (**D**) Quantification results of cell aggregation in PBS. The proportion of aggregated Raji cells was calculated as PE-FITC double positive/PE positive. (**E**) The blocking effect of human serum IgG on the FcγRCARs. JK-FcγRCAR cells were incubated in human serum, PBS with 10% human serum, or PBS for 30 min. The captured IgG on JK-FcγRCAR cells was detected with goat anti-human IgG antibody (APC). (**F**) The fluorescence microscope detection of cell aggregation after a 3 h of co-incubation in human serum. (**G**) Quantification results of cell aggregation in human serum. Data are presented as mean ± SD, n = 3. Statistical analysis was based on two-tailed heteroscedastic Student’s t-test, and statistical significance is considered at p<0.05. When differences are statistically significant, the significance is represented with asterisks (*) according to the following values: p<0.05 (*), p<0.01 (**), and p<0.001 (***). “n.s.” indicates not significant (p>0.05).

The effector and target cells’ aggregation was quite apparent in the groups of Raji-Rtx cells & JK-FcγRCAR cells but not in the others (Fig. 3B). According to the results of FACS, the ratios of aggregated Raji cells (calculated as FITC^+^PE^+^/PE^+^) were 16.3±2.51%, 29.06±2.73%, and 58.09±1.14% in Raji-Rtx cells & JK-16CAR, JK-32CAR, and JK-64CAR cells, respectively (Fig. 3C and 3D). In contrast, there were less than 8% doublets without Rtx, or with JK-EGCAR cells regardless of whether Rtx was present. These results show that the affinity play a key effect on the cell aggregation.

### 3.4 Blockage of serum IgG on FcγRCARs

We then investigated the block effect of high-level serum IgG (~10 mg/mL) on FcγRCARs, and found significant enhancement of APC fluorescence in all the three FcγRCAR groups, which demonstrated their binding to serum IgG (Fig. 3E). Almost the same APC signal was detected when the human serum’s concentration was decreased to 10%, suggesting that nearly all the FcγRCARs were blocked in 10% human serum. To further explore their activities in the human body, JK-FcγRCAR-DiO cells and Raji-DiI cells pre-incubated with Rtx or not were resuspended in human serum. After a 3 h co-incubation, FcγRCARs mediated aggregation were observed for Raji-DiI-Rtx cells, but not Raji-DiI cells (Fig. 3F). The ratios of aggregated Raji cells in PBS and human serum were close for 16CAR (16.3±2.51% vs. 17.08±1.47%) and 32CAR (29.06±2.7% vs. 26.5±2.62%), while a moderate decrease was observed for 64CAR (58.09±1.15% vs. 49.44±2.82%) (Fig. 3G).

### 3.5 Formation of antibody-antigen immune complex induces JK-FcγRCAR cells activities

The cytotoxic activity of JK-FcγRCAR cells was assessed with Raji cells and BT474 cells (Her2^+^) as tumor targets, with Rtx and Hct as the adaptors, respectively. Firstly, the high-level expression of the two antigens was confirmed by FACS that the antigens on 1E4 Raji cells or 3E3 BT474 cells in 100 μL could be saturated by 0.1 μg/mL Rtx or Hct (Fig. S3). Given that the FcγRCARs would also be saturated by 0.1 μg/mL antibody, the *in vitro* tumor killing assay concentrations were set between 0-0.1 μg/mL, and the E: T ratio was set at 5. In the 4 h LDH release cytotoxicity assay, the JK-FcγRCAR cells displayed an ADCC effect which heavily depends on both the concentration of antibodies and the affinity of the FcγRCARs (Fig. 4A and 4B). Target cell killing was low in the absence of the antibodies or with JK-EGCAR cells.

**Figure 4.**
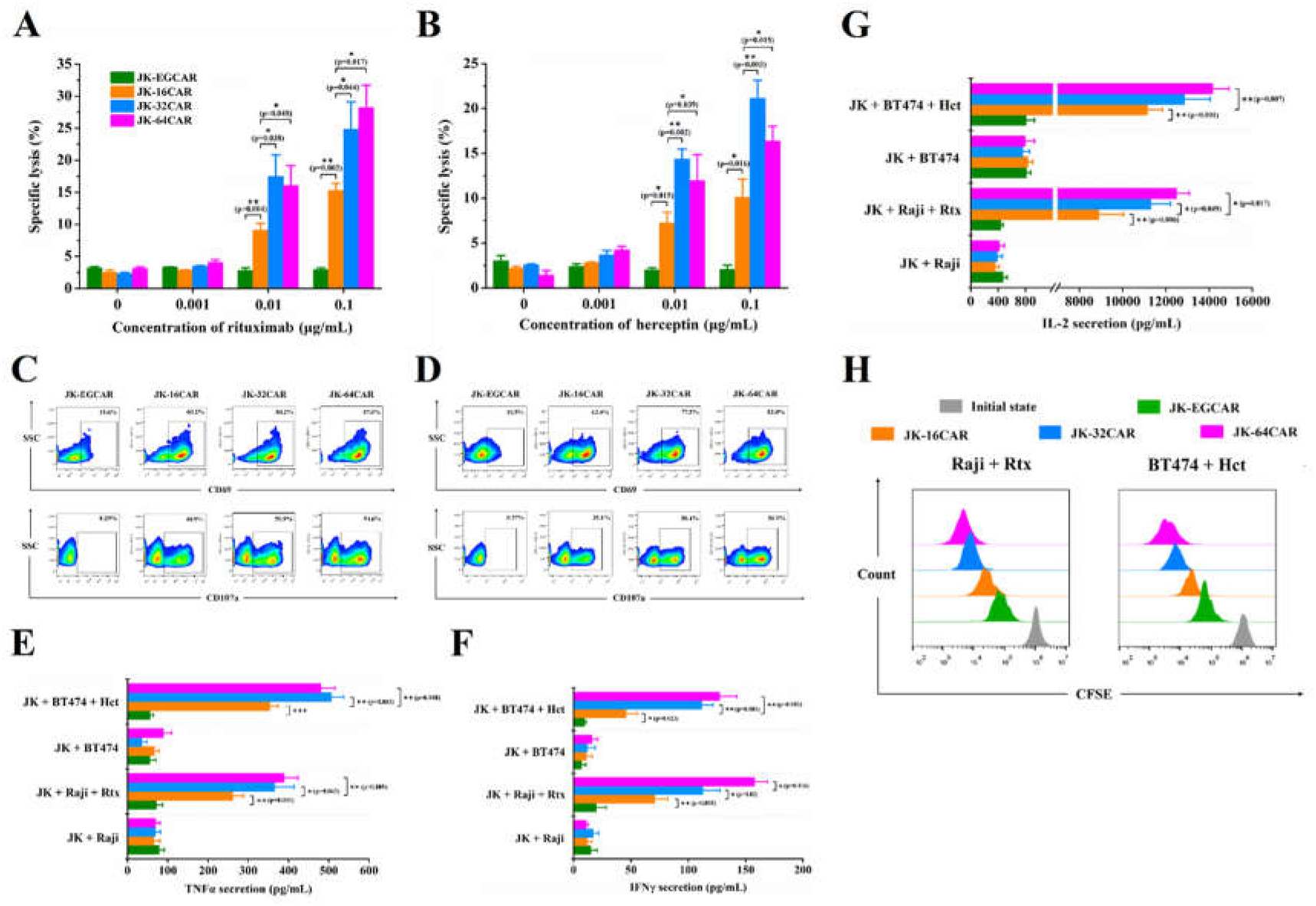
ADCC, cell activation, cytokine release, and cell proliferation effects mediated by immune complexes. The specific lysis for rituximab opsonized Raji cells (**A**), and herceptin opsonized BT474 cells (**B**) at various concentrations of antibodies detected by LDH Cytotoxicity Assay Kit after 4 h of co-incubation (E: T ratio=5). (**C**& **D**) The up-regulation of CD69 and CD107a of JK-EGCAR/FcγRCAR cells at 0.1 μg/mL antibodies. TNFα (**E**), IFNγ (**F**), and IL-2 (**G**) secretion with 10% FBS and 48 h of co-incubation, measured using ELISA kits. (**H**) The cell proliferation of JK-EGCAR/FcγRCAR cells with 10% FBS and 24 h of co-incubation. Data are presented as mean ± SD, n = 3. Statistical analysis was based on two-tailed heteroscedastic Student’s t-test, and statistical significance is considered at p<0.05. When differences are statistically significant, the significance is represented with asterisks (*) according to the following values: p<0.05 (*), p<0.01 (**), and p<0.001 (***). “n.s.” indicates not significant (p>0.05).

We further evaluated whether FcγRCARs cross-linking by immune complex could induce activation signals and exocytosis of lytic granules, cytokines secretion, and cell proliferation. After 4 h of co-culture, JK-FcγRCAR cells, but not JK-EGCAR cells, up-regulated the expression of CD69 and CD107a markedly as a result of recognizing Rtx opsonized Raji cells (Fig. 4C) or Hct opsonized BT474 cells (Fig. 4D). During the 48 h of the co-culture process, JK-FcγRCAR cells produced much higher levels of TNFα (Fig. 4E), IFNγ (Fig. 4F), and IL-2 (Fig. 4G) than JK-EGCAR cells in response to the immune complexes, but not sole antibodies (data not shown). Finally, the CFSE dilution assay demonstrates a more robust proliferative response mediated by FcγRCARs with higher affinity (Fig. 4H). Apparent correlations between the receptors’ affinities and the intensities of all these cellular outputs were observed.

### 3.6 De-glycosylated antibodies maintain binding to 64CAR, but not the other FcγRCARs

To examine the binding propensities of the FcγRCARs to de-glycosylated antibodies, Rtx was digested by PNGase F for 24 h, and the complete removal of the attached sugar was confirmed by reducing SDS-PAGE (Fig. S3). In the FACS analysis, no detectable binding to either 16CAR or 32CAR could be measured for de-glycosylated Rtx (Dg-Rtx), but 64CAR maintained some extent of binding (Fig. 5A). To characterize the cell activating and ADCC effects such a binding capacity can mediate, cell aggregation tests were performed with Rtx replaced by de-glycosylated Rtx (Dg-Rtx). The binding ratio of Raji cells by JK-64CAR cells was decreased to 30.45±1.7%, whereas the binding of 16CAR and 32CAR to Fc fragment were totally abolished (Fig. 5B). The results of microscopic fluorescence inspection provided further confirmation (Fig. 5C). At 0.1 μg/mL, Dg-Rtx mediated a 21.27±2.91% Raji cells killing effect by JK-64CAR cells, and this observation was reproducible using BT474 and Dg-Hct (13.36±2.33% killing effect) (Fig. 5D). It appears that the combination of JK-64CAR cells and de-glycosylated antibody exhibited comparable target lysing ability to the combination of JK-32CAR cells and primary antibody, owing to the similar affinities.

**Figure 5.**
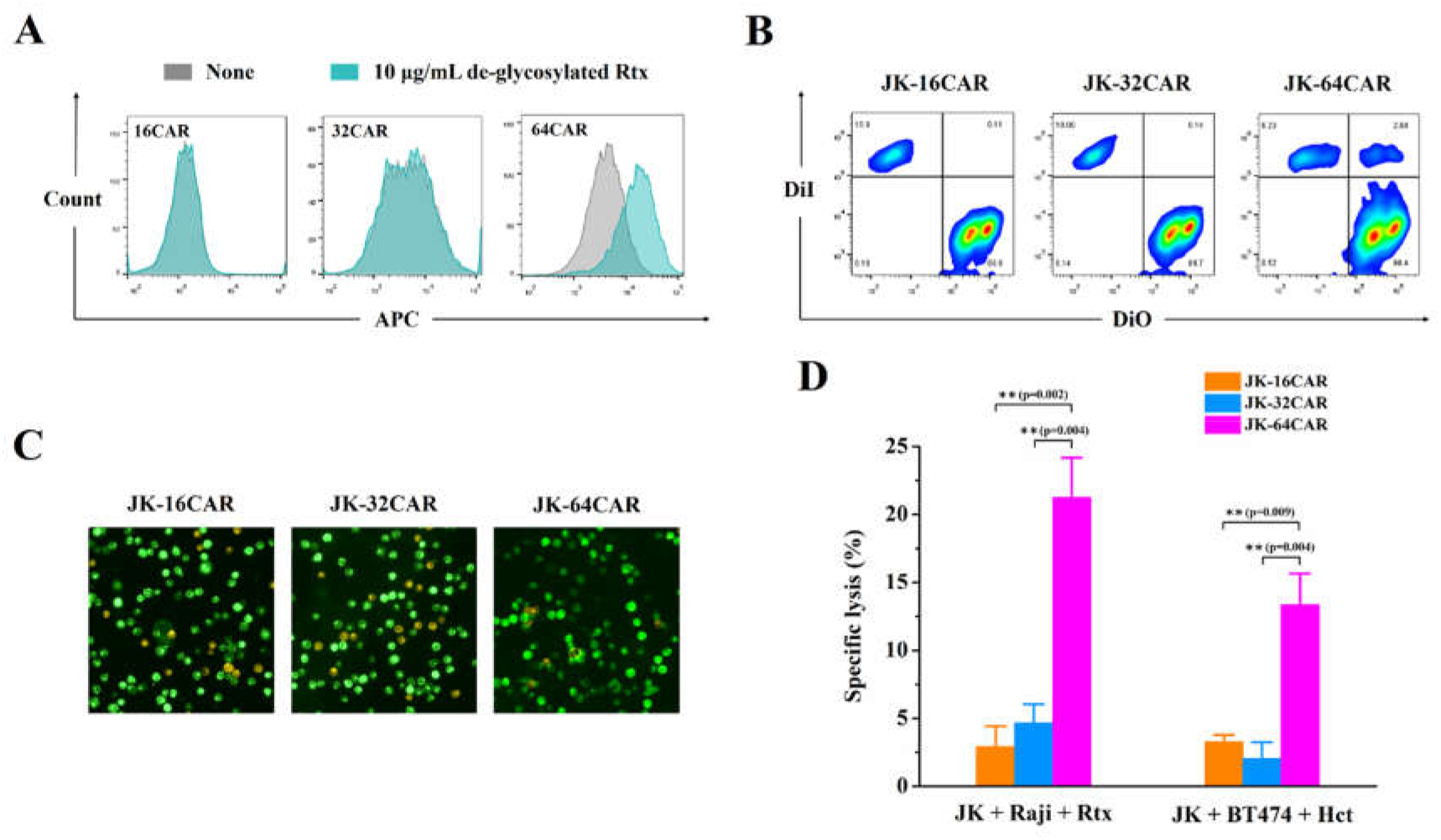
64CAR’s binding to the de-glycosylated antibody. (**A**) FACS analysis of the bound de-glycosylated rituximab on the FcγRCARs. JK-FcγRCAR cells were incubated with 10 μg/mL rituximab and goat anti-human IgG antibody (APC) in sequence, and then cell staining was measured by FACS. (**B**) FACS analysis of the ratio of aggregated Raji cells after 1 h co-incubation with JK-FcγRCAR cells (E: T ratio=10) and 0.1 μg/mL de-glycosylated rituximab in PBS. The aggregated Raji cells were shown as the PE-FITC double-positive events. (**C**) The detection of cell aggregation with a fluorescence microscope. (**D**) The specific lysis for de-glycosylated rituximab opsonized Raji cells and de-glycosylated herceptin opsonized BT474 cells detected by LDH Cytotoxicity Assay Kit after 4 h of co-incubation (E: T ratio=5). Data are presented as mean ± SD, n = 3. Statistical analysis was based on two-tailed heteroscedastic Student’s t-test, and statistical significance is considered at p<0.05. When differences are statistically significant, the significance is represented with asterisks (*) according to the following values: p<0.05 (*), p<0.01 (**), and p<0.001 (***). “n.s.” indicates not significant (p>0.05).

### 3.7 Comparison of the three FcγRCARs in the tumor model with low antigen density

*In vivo* antitumor activity mediated by combined administration of JK-FcγRCAR cells and IgG1 antibody was measured using xenografted nude mouse models (Fig. 6A). On day0, 5E6 U251 MG cells (with Matrigel) were inoculated subcutaneously in the right flank of female nude mice. Palpable solid tumor xenografts (~200 mm^3^) were generated 8 days after tumor implantation, and the expression of Her2 on the surface of the transplanted tumor cells was verified by FACS (Fig. S5). Significant tumor inhibition (Fig. 6B–6D) and increased Jurkat cells’ proliferation in tumor tissue (Fig. 6E, Fig. S6) were only observed in the JK-64CAR & Hct group. In contrast, no significant difference in cell proliferation was observed in peripheral blood (Fig. 7A, Fig. S7) or spleen (Fig. 7B, Fig. S8), or tumor tissue in the absence of Hct. These results indicate that Fcγ receptors can mediate effector cells to recognize and kill tumor cells in an immune complex-dependent manner strictly. Higher levels of intratumoral hTNFα and hIL-2 also demonstrate the more robust activation of JK-64CAR cells (Fig. 7C). Besides, the more vigorous proliferation of the endogenous lymphocytes in tumor tissue was detected in the JK-64CAR cells & Hct group (Fig. 7D, Fig. S6), suggesting a positive cross-talk between the JK-64CAR cells and the endogenous immune system. Altogether, the JK-64CAR cells exhibited more potent antitumor activity without causing significant side effects (Fig. 7E) in the low Her2 density U251 MG model compared with the lower-affinity counterparts.

**Figure 6.**
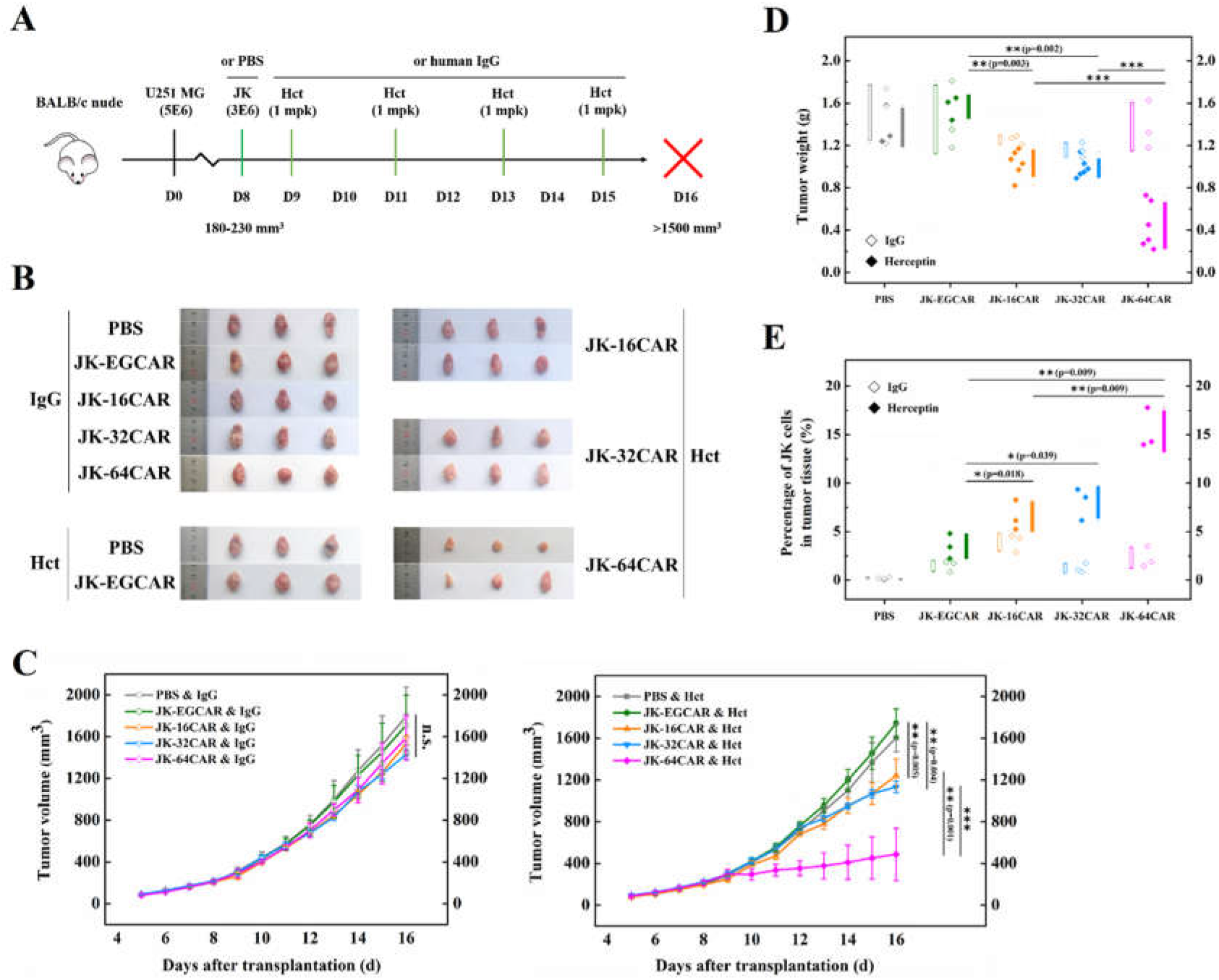
Efficacy of combination treatment of JK-FcγRCAR cells and herceptin in the U251 MG xenograft model. (**A**) Schematic depiction of tumor inoculation and treatment protocol. Female nude mice were implanted s.c. with 5E6 U251 MG cells on day0. On day8, 3E6 JK-EGCAR/FcγRCAR cells or PBS were injected i.v. into mice. From day9, mice were injected i.v. with herceptin or human IgG (1 mg/kg) every other day and executed on day16. (**B**) Digital image of the stripped tumors. (**C**) Tumor volume of the ten groups. (**D**) Tumor weight of the ten groups. (**E**) On day16, the percentage of JK-EGCAR/FcγRCAR cells in tumor tissue was detected by FACS. Each point represents one mouse. Data are presented as mean ± SD, n = 3 for control groups and n = 6 for experimental groups. Statistical analysis was based on two-tailed heteroscedastic Student’s t-test, and statistical significance is considered at p<0.05. When differences are statistically significant, the significance is represented with asterisks (*) according to the following values: p<0.05 (*), p<0.01 (**), and p<0.001 (***). “n.s.” indicates not significant (p>0.05).

**Figure 7.**
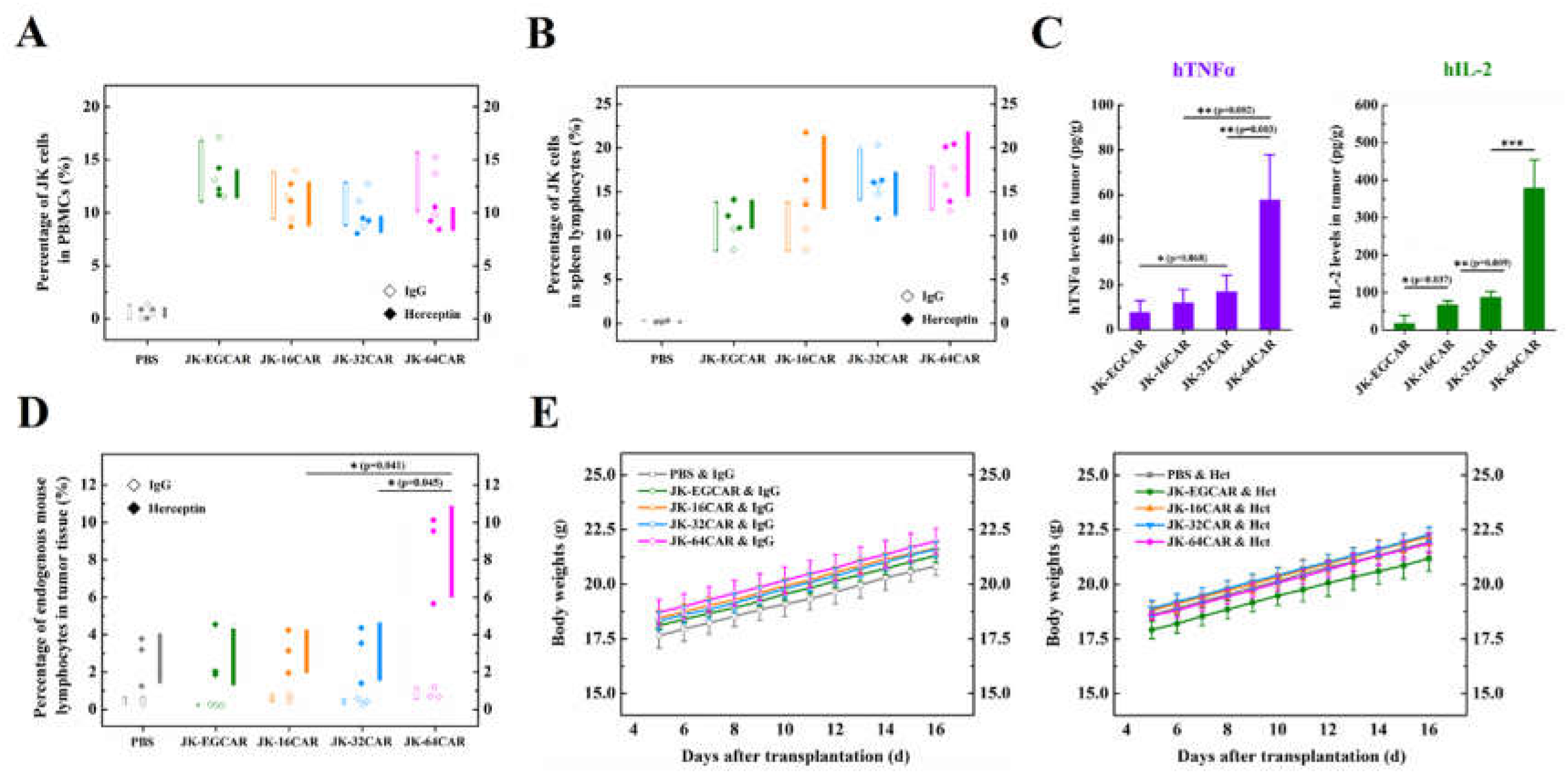
Analysis of the therapeutic and side effect profiles in the U251 MG model. On day16, the percentage of JK-EGCAR/FcγRCAR cells in peripheral blood (**A**) and spleen lymphocytes (**B**) was detected by FACS. (**C**) Levels of hTNFα and hIL-2 in tumor tissue, detected by ELISA kits. (**D**) FACS analysis of the percentage of endogenous mouse lymphocytes in tumor tissue. (**E**) Body weight of the ten groups. Each point represents one mouse. Data are presented as mean ± SD, n = 3 for control groups and n = 6 for experimental groups. Statistical analysis was based on two-tailed heteroscedastic Student’s t-test, and statistical significance is considered at p<0.05. When differences are statistically significant, the significance is represented with asterisks (*) according to the following values: p<0.05 (*), p<0.01 (**), and p<0.001 (***). “n.s.” indicates not significant (p>0.05).

### 3.8 Comparison of the three FcγRCARs in the tumor model with high antigen density

To investigate the *in vivo* bioactivities of the JK-FcγRCAR cells in the tumor model with high antigen density, we subcutaneously inoculated 4E6 Raji cells in the right flank of female nude mice (Fig. 8A). Palpable solid tumor xenografts (~200 mm^3^) were generated 10 days after tumor implantation, and the high expression of CD20 on the transplanted tumor cells was verified by FACS (Fig. S9). As shown in Fig. 8B–8D, co-administered JK-FcγRCAR cells and Rtx obviously suppressed the tumor growth during the 8-day observation period compared with control groups in which Rtx was replaced by human IgG or JK-FcγRCAR cells were replaced by JK-EGCAR cells or PBS. In line with their higher affinity for Fc, 32CAR and 64CAR induced significantly stronger tumor inhibition effect and increased JK-FcγRCAR cells’ proliferation in tumor tissue (Fig. 8E, Fig. S10), but not in peripheral blood (Fig. 8F, Fig. S11) or spleen (Fig. 8G, Fig. S12), or tumor tissue in the absence of Rtx. Consistent with the results of the *in vitro* experiments, positive correlations between affinity and effect strengths were observed at the initial stage of therapy (Fig. 8C). However, mice treated with JK-64CAR cells & Rtx developed symptoms of systemic toxicity indicated by weight loss (Fig. 9A), hunched bodies, and motor weakness later. Though also found to an obviously lesser extent, these symptoms went away quickly and did not affect the durable efficacy in the JK-32CAR cells & Rtx group. We further examined the immunological status of tumor infiltrated JK-FcγRCARs cells by FACS. As shown in Fig. 9B and Fig. S13, the expression of PD-1 increased along with the increase of the affinity of the FcγRCAR, whereas the expression of CD25 was down-regulated in the JK-64CAR cells & Rtx group. As a CD4^+^ T cell line, Jurkat cells could modulate the tumor immune environment. The more vigorous proliferation of infiltrating endogenous mouse lymphocytes was also detected in the JK-32CAR cells & Rtx group (Fig. 9C, Fig. S10), implying an activation-prone immune contexture. Besides higher plasma levels of hTNFα and hIL-2 (Fig. 9D), 64CAR significantly elevated plasma mIL-6 and mIL-10. Such pro-inflammatory and immunosuppressive cytokine profiles probably adversely affected the antitumor responses. Overall, the superior antitumor efficacy and safety profiles of JK-32CAR cells point out that a universal CAR with 10^−7~8^ M affinity can retain an optimal effector and safety in a solid tumor with high antigen density.

**Figure 8.**
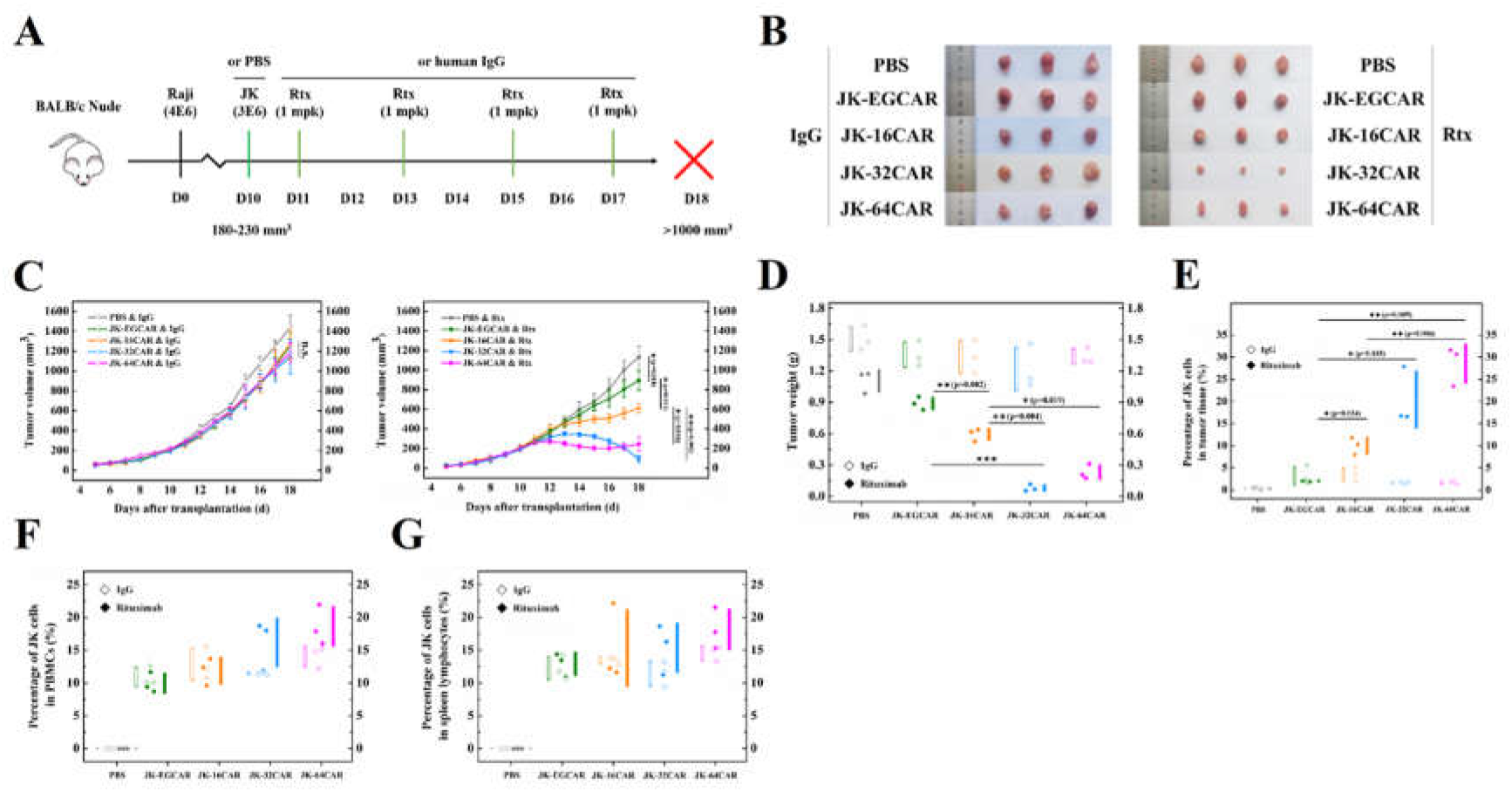
Efficacy of combination treatment of JK-FcγRCAR cells and rituximab in the Raji xenograft model. (**A**) Schematic depiction of tumor inoculation and treatment protocol. Female nude mice were implanted s.c. with 4E6 Raji cells on day0. On day10, 3E6 JK-EGCAR/FcγRCAR cells or PBS were injected i.v. into mice. From day11, mice were injected i.v. with rituximab or human IgG (1 mg/kg) every other day and executed on day18. (**B**) Digital image of the stripped tumors. (**C**) Tumor volume of the ten groups. (**D**) Tumor weight of the ten groups. On day18, the percentage of JK-EGCAR/FcγRCAR cells in tumor tissue (**E**), peripheral blood (**F**), and spleen lymphocytes (**G**) were detected by FACS. Each point represents one mouse. Data are presented as mean ± SD, n = 3. Statistical analysis was based on two-tailed heteroscedastic Student’s t-test, and statistical significance is considered at p<0.05. When differences are statistically significant, the significance is represented with asterisks (*) according to the following values: p<0.05 (*), p<0.01 (**), and p<0.001 (***). “n.s.” indicates not significant (p>0.05).

**Figure 9.**
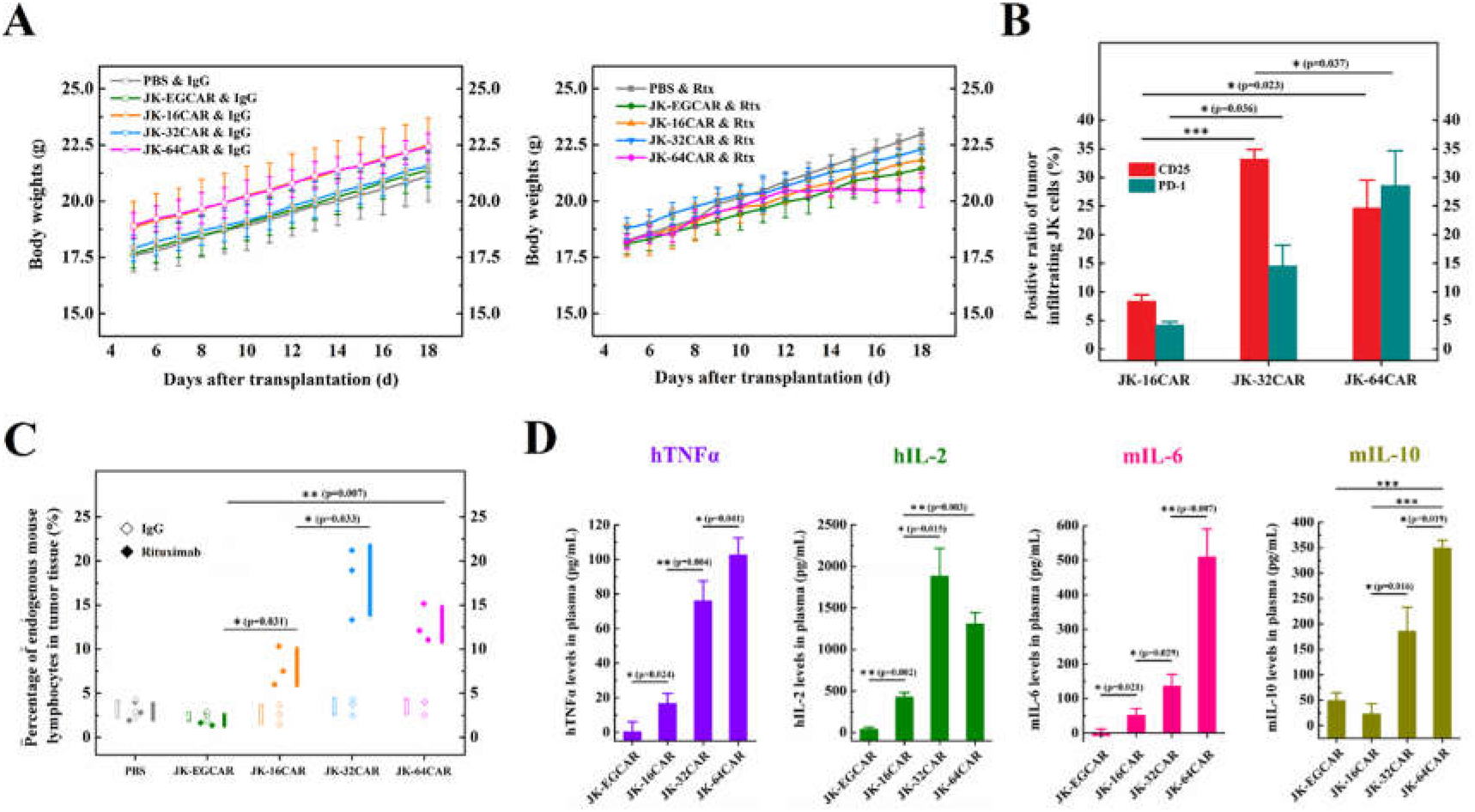
Analysis of side effects, immunosuppression, and cytokines profiles in the Raji xenograft model. (**A**) Body weight of the ten groups. (**B**) FACS analysis of the CD25 and PD-1 expression on tumor-infiltrating JK-FcγRCAR cells. (**C**) FACS analysis of the percentage of endogenous mouse lymphocytes in tumor tissue. (**D**) Levels of hTNFα, hIL-2, mIL-6, and mIL-10 in mouse plasma, detected by ELISA kits. Each point represents one mouse. Data are presented as mean ± SD, n = 3. Statistical analysis was based on two-tailed heteroscedastic Student’s t-test, and statistical significance is considered at p<0.05. When differences are statistically significant, the significance is represented with asterisks (*) according to the following values: p<0.05 (*), p<0.01 (**), and p<0.001 (***). “n.s.” indicates not significant (p>0.05).

### 3.9 Influence of antibody-antigen affinity on the in vivo bioactivities of JK-FcγRCAR cells

To further explore the influence of antibody-antigen affinity on the *in vivo* bioactivities of JK-FcγRCAR cells, we performed N93K site mutation in the light chain and D57E site mutation in the heavy chain of Rtx, respectively, to obtain Rtx variants with different affinities (Fig. 10A). The Rtx-M produced in our lab bound to CD20 with a high affinity (4.17 nM), comparable with commercial rituximab (3.66 nM) (Fig. 10B). The affinities of Rtx-L (12.51 nM) and Rtx-H (0.93 nM) to CD20 were also consistent with the previous report^30^. The antitumor activities of JK-FcγRCAR cells & Rtx-L/M/H were evaluated in the Raji model (Fig. 10C). We found that the influence of antibody-antigen affinity was only observed in the JK-16CAR group (Fig. 10D-10F). In the group of Rtx with higher affinity, the JK-16CAR cells exhibited more robust proliferation (Fig. 10G, Fig. S14) and produced a larger amount of hIL-2 (Fig. 10H) in tumor tissue. For the low-affinity 16CAR, it is beneficial to combine it with a higher-affinity antibody.

**Figure 10.**
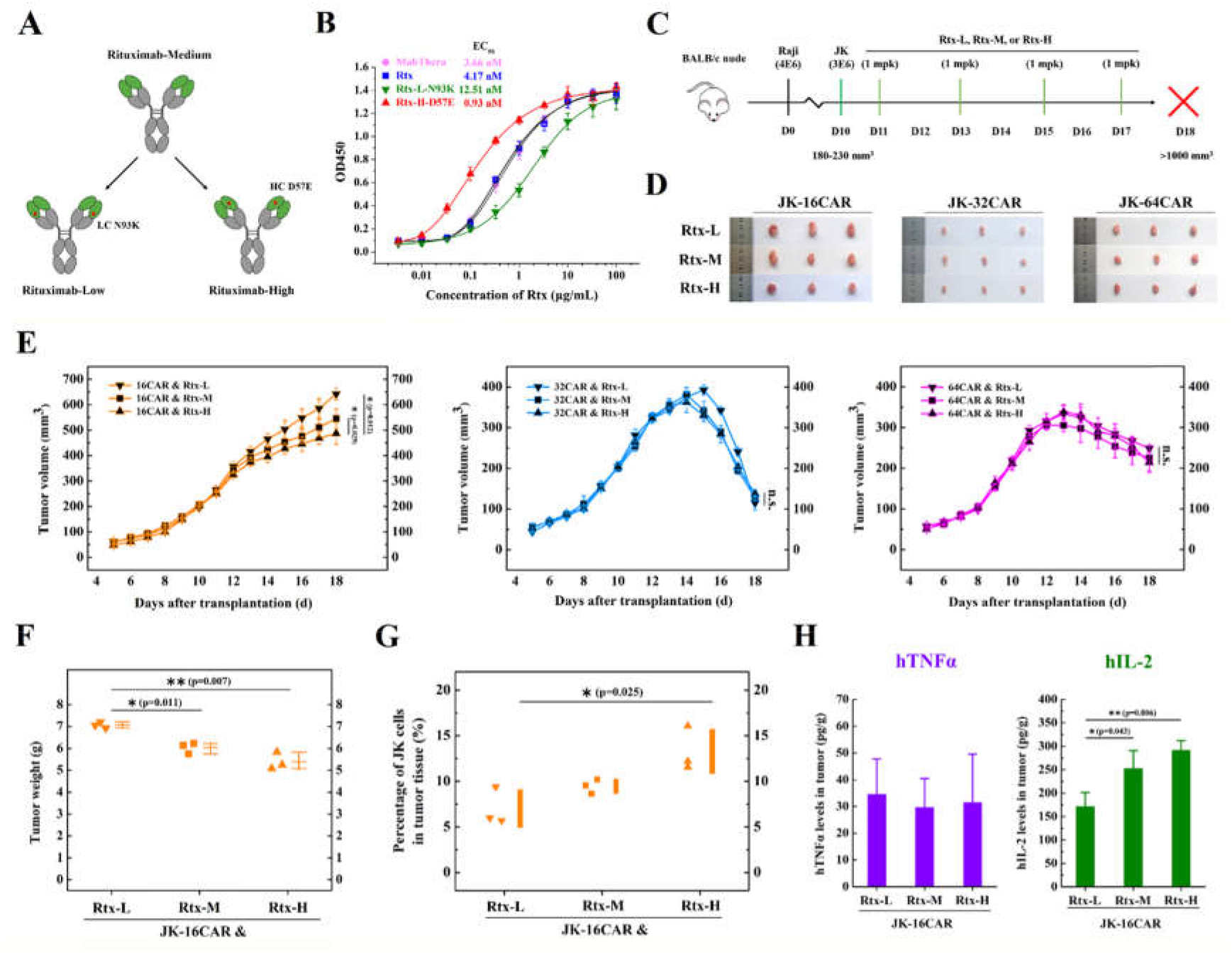
Analysis of the influence of antibody affinity in the Raji xenograft model. (**A**) Schematic depiction of site mutations on rituximab. (**B**) Affinity detection of rituximab and its variants by ELISA. (**C**) Schematic depiction of tumor inoculation and treatment protocol. Female nude mice were implanted s.c. with 4E6 Raji cells on day0. On day10, 3E6 JK-FcγRCAR cells were injected i.v. into mice. From day11, mice were injected i.v. with rituximab-L/M/H (1 mg/kg) every other day and executed on day18. (**D**) Digital image of the stripped tumors. (**E**) Tumor volume of the nine groups. (**F**) Tumor weight of the JK-16CAR & Rxt-L/M-H groups. (**G**) On day18, the percentage of JK-16CAR cells in tumor tissue was detected by FACS. (**H**) Levels of hTNFα and hIL-2 in tumor tissue, detected by ELISA kits. Each point represents one mouse. Data are presented as mean ± SD, n = 3. Statistical analysis was based on two-tailed heteroscedastic Student’s t-test, and statistical significance is considered at p<0.05. When differences are statistically significant, the significance is represented with asterisks (*) according to the following values: p<0.05 (*), p<0.01 (**), and p<0.001 (***). “n.s.” indicates not significant (p>0.05).

## 4. Discussion

Though CAR T cells have achieved significant success in treating several hematological malignancies, the applicability of this therapy urgently needs to be expanded. Emerging approaches based on combinations of universal CAR T cells and their respective adaptor molecules, which mediate the crosslinking between CAR T effector cells and target, have been developed. For the scFv-based universal CAR, the unpredictable expression level and tonic signaling caused by self-aggregation in the absence of antigen are tricky problems^32^. Besides, the conformation of the antibody was influenced by the tagged small PNE (14-aa), and the tagging site determined the distance and orientation between effector cells and target, thus affecting recognition and cytotoxicity^9^. Last but not least, until now, no clinical data reported the immunogenicity of these tags, and eventual death of mice due to the emergence of anti-FITC immunity was observed^8^. Several groups tried to overcome these drawbacks by engineering T cells to express a CAR where the scFv is replaced by the outer membrane domain of the FcγRIIIA (CD16a) and utilizing the clinically approved IgG1 mAbs function as the adaptors^21–29^. When infused concomitantly with mAb, CD16a-T cells, but not rhIL-2–treated NK cells, significantly suppressed the tumor growth in mice, which demonstrated the promising potential of the CD16a-based universal CAR T-cell therapy^24^.

Although targeting various antigens, Mabs’ engagement with Fc receptors on multitudinous effector cells is almost universal, thus making themselves natural adaptors in the human body. Numerous techniques have been employed to improve Fc functions. Defucosylation of the glycan moiety attached to IgG-Fc^33^ or site mutations^34^ can significantly increase the binding affinity for CD16a, thus generating a remarkable enhancement of ADCC responses. On the other hand, FcγR polymorphisms also influence patients’ response to treatment with therapeutic mAbs. Allotypes of CD16a result in receptors with either a phenylalanine (F) or a valine (V) residue at position 158; CD16a V158, which occurs in a minority of individuals, has stronger Fc binding and is associated with superior clinical responses^35^. In line with these, Felicitas et al. proved that the high-affinity V158 variant renders CD16a-CAR T cells superior to the more prevalent F158 low-affinity variant^29^. These highlight the significant meaning to optimize the affinity of FcγRCARs.

To develop an affinity fine-tuned universal CAR, here we performed a side-by-side investigation of FcγRIIIA-F (CD16a-F), FcγRIIA-H (CD32a-H), and FcγRI (CD64), which possess different affinities for antibodies with KD values between 10^−6^ M and 10^−9^ M, for their potential as chimeric receptors. *In vitro* experiments show that the bio-activities of JK-FcγRCAR cells, including cell activation, proliferation, cytokines secretion, and ADCC effect, were entirely dependent on both the FcγRCARs and the antibody. Even though the differences between 32CAR and 64CAR were not significant, positive correlations between the affinity and the effect strength were observed. Furthermore, within the clinically relevant concentrations of 0-0.1 μg/mL, we proposed that the efficacy and off-target effect can be regulated by adjusting the antibody titer in plasma.

It has been widely shown that deglycosylation of antibodies resulted in a loss of effector function—an inability to bind CD16a and induce ADCC effect^36^. In consistence with the previous work, the de-glycosylated antibody failed to bind all FcγRCARs except 64CAR^37^. Accompanied with the finding that de-glycosylated IgG displayed similar antigen binding, solubility, stability at physiological temperature, and serum persistence *in vivo* to those of glycosylated IgG^38^, our results provide support for the development of de-glycosylated antibodies for clinical trials.

We then explored the antitumor activities of the three FcγRCAR T cells in mice. In the Her2 low expression U251 MG model, only the JK-64CAR cells & Hct combination displayed considerable tumor inhibition effect, indicating the crucial role of high affinity. Probably, the higher affinity of 64CAR facilitates more efficient recognition of the JK-64CAR cells to the low-density antigen. In the CD20 high expression Raji model, JK-FcγRCAR cells exhibited antigen-dependent and antibody-dependent effects in the presence of Rtx. There was no significant difference between the three FcγRCARs groups in the percentage of effector cells in splenic and peripheral between different groups, indicating that the FcγRCARs redirected effector cells without the need for major histocompatibility complex (MHC) restriction, similar to the conventional CARs. At a low dosage (1 mg/kg every other day) of Rtx, the affinity of FcγRCAR seems to be a crucial determinant for the outcome of this form of therapy. In combination with Rtx, JK-16CAR cells displayed weak activation, proliferation, and tumor growth inhibition. Comparatively, 32CAR and 64CAR mediated much more potent effects, with severe CRS and immunosuppression in the JK-64CAR cells & Rtx group.

Compared with antibody monotherapy, the high affinity and strong activation effects of CARs bring not only excellent therapeutic effects but also much more sensitive immune responses. Over-activation of CAR T cells easily causes even lethal CRS and immune effector cell-associated neurotoxicity syndrome (ICANS) in patients^39^. The core cytokines in CRS, such as IL-6, IL-10, and TNFα, also support heavily immunosuppressive context, thus transferring the effector cells to states of dysfunction, including anergy, exhaustion, and senescence^40, 41^. A lower-affinity CD19 CAR (KD=14 nM) showed enhanced antitumor activity, more excellent persistence, and better cytokine phenotype in patients, than the CAR derived from FMC63, the high-affinity antibody (KD=0.3 nM) used in many clinical studies^42^. Other works also reported the benefits of using moderate-affinity CAR ranging from 10-100 μM^20^. Silvia et al. demonstrate that a 10 μM binding affinity for the target antigen was sufficient to ensure robust functional responses^13^. We assumed that CAR functionality could be delicately modulated by changing the binding affinity to the cognate antigen.

Combined with the JK-FcγRCAR cells, rituximab variants with different affinities exhibited similar *in vivo* antitumor efficacy. We presume that the antibody-antigen affinity primarily determines antibody distribution and show a saturation effect. In a study examining the effects of affinity on *in vivo* tumor targeting of herceptin, tumor targeting capacity would not be enhanced once its KD value reached 10 nM^43^. Lower affinity antibodies even penetrated farther into the tumor tissue due to their ability to dissociate from antigen following binding and continue diffusing through the tumor interstitial space^44^. Unlike the affinity between FcγR and antibody, the affinity between antibody and antigen seems to have a minor impact on the bioactivity of the effector cells.

A big concern about FcγRCAR T cells is that the therapeutic mAbs may compete with high-level serum immunoglobulins (~10 mg/mL) for their binding to the FcγRs and may be defective in their ability to mediate ADCC. Although it is widely accepted that CD16a and CD32a have no meaningful binding to monomeric IgG for their low affinities, the current study’s findings do not support this idea. In this study, 16CAR and 32CAR displayed detectable binding to 1 μg/mL IgG in antibody binding assay and can be saturated in 10% human serum (1 mg/mL IgG). Results of the binding assay in another study indicate that at physiological concentrations, IgG would occupy 98.5% and 97.4% of CD32a and CD16a, respectively^45^. On the other hand, we determined that the pre-bound monomeric IgG on FcγRCARs would be substituted by the immune complex, which several mechanisms can explain. Binding to antigen induces conformational changes of antibody from “close” to “open” state, making it easier to be captured by FcγRs^46^. Multivalent cross-linking of FcγRs with immune complexes provided stronger binding. It has been shown that inside-out signaling could contribute to immune complex binding to FcγRs occupied by serum IgG^47^. Stimulation with cytokines, such as TNFα and IFNγ, increased the binding of CD64 to immune complexes by sophisticated cytoskeleton rearrangements^48^. While detecting an enhanced activity mediated by CD16a-CAR in the presence of high doses of polyclonal IgGs, Felicitas et al. attributed it to the unspecific deposition of IgGs on tumor cells. They noted that doses required for polyclonal triggering of CD16a-CAR T cells were several log-concentrations higher than for mAbs^29^. In a word, the binding of serum IgG on the FcγRCARs may not cause safety concerns.

Our research has several limitations. Although Jurkat cells have been extensively used as the model to characterize signaling events in T cell activation, they differ from primary T cells in many ways. Jurkat cells chimeric with FcγRCARs secreted a large amount of IL-2, but little TNFα and IFNγ, accompanied with disappointing *in vitro* ADCC effects. The second is that the on-target off-tumor side effects were not fully explored since Rtx/Hct neither bind mouse CD20/Her2 nor normal tissues of mice. The humanized mouse model, or antibodies that can react with both human and mouse antigens, may provide solutions. Lastly, further work is required to investigate the *in vivo* influence of human serum IgG on the efficacy and safety of FcγRCARs. All of these are important issues for future research.

Although further studies are warranted, our research strongly suggests the significance of utilizing Fc receptors with a moderate affinity for antibodies to develop universal CARs. Our work underlines the great potential of FcγRCAR T cells and antibody combination therapy, in which the specific recognition of antigens by Fab, the bridge function of Fc and FcγR, and the powerful tumor-killing capacity of T cells would be finely combined.

## 5. Conclusion

In summary, the CARs chimeric with three different affinity Fcγ receptors, namely CD16a, CD32a, and CD64, were compared side by side. The activities of these universal CAR T cells toward tumor cells, which include cell activation, proliferation, cytokines secretion, and ADCC effect, can be redirected and regulated by IgG1 antibodies. Positive correlations between the affinities of the FcγRCARs and the intensity of these effects were observed *in vitro*. While significantly increasing the tumor inhibition effect in the Her2 low expression U251 MG model, 64CAR’s strong affinity led to over-activation of effector cells and subsequent CRS, immunosuppression, and tumor return in the CD20 high expression Raji model. Comparatively, the moderate affinity of 32CAR achieved a better balance between efficacy and side effects. Altogether, our study explored the Fcγ receptors-based universal CAR T therapy and demonstrated the profound influence of affinity.

## Abbreviations

ADCC: antibody-dependent cell-mediated cytotoxicity
AICD: activation-induced cell death
CAR: chimeric antigen receptor
CFSE: carboxyfluorescein succinimidyl ester
CRS: cytokine release syndrome
ELISA: enzyme-linked immunosorbent assay
FACS: fluorescence-activated cell sorting
FBS: fetal bovine serum
FcγR: Fcγ receptor
Hct: herceptin
IgG: immunoglobulin G
LDH: lactate dehydrogenase
mAb: monoclonal antibody
Rtx: rituximab
scFv: single chain antibody fragment
SD: standard deviation
TAA: tumor-associated antigen

## References

1. Ying Z, Huang XF, Xiang X, Liu Y, Kang X, Song Y, et al. A safe and potent anti-CD19 CAR T cell therapy. Nat Med 2019; 6:947–953. http://dio.org/10.1038/s41591-019-0421-7.

2. Benmebarek MR, Karches CH, Cadilha BL, Lesch S, Endres S, Kobold S. Killing Mechanisms of Chimeric Antigen Receptor (CAR) T Cells. Int J Mol Sci 2019; 20:1283. http://doi.org/10.3390/ijms20061283.

3. Marofi F, Motavalli R, Safonov VA, Thangavelu L, Yumashev AV, Alexander M, et al. CAR T cells in solid tumors: challenges and opportunities. Stem Cell Res Ther 2021; 12:81. http://doi.org/10.1186/s13287-020-02128-1.

4. Wagner J, Wickman E, DeRenzo C, Gottschalk S. CAR T Cell Therapy for Solid Tumors: Bright Future or Dark Reality? Mol Ther 2020; 28: 2320–2339. http://doi.org/10.1016/j.ymthe.2020.09.015.

5. Martinez M, Moon EK. CAR T Cells for Solid Tumors: New Strategies for Finding, Infiltrating, and Surviving in the Tumor Microenvironment. Front Immunol 2019; 10:128. http://doi.org/10.3389/fimmu.2019.00128.

6. Poorebrahim M, Sadeghi S, Fakhr E, Abazari MF, Poortahmasebi V, Kheirollahi A, et al. Production of CAR T-cells by GMP-grade lentiviral vectors: latest advances and future prospects. Crit Rev Clin Lab Sci 2019; 56:393–419. http://doi.org/10.1080/10408363.2019.1633512.

7. Darowski D, Kobold S, Jost C, Klein C. Combining the best of two worlds: highly flexible chimeric antigen receptor adaptor molecules (CAR-adaptors) for the recruitment of chimeric antigen receptor T cells. MAbs 2019; 11:621–631. http://doi.org/10.1080/19420862.2019.1596511.

8. Tamada K, Geng D, Sakoda Y, Bansal N, Srivastava R, Li Z, Davila E. Redirecting gene-modified T cells toward various cancer types using tagged antibodies. Clin Cancer Res 2012; 18:6436–6445. http://doi.org/10.1158/1078-0432.CCR-12-1449.

9. Rodgers DT, Mazagova M, Hampton EN, Cao Y, Ramadoss NS, Hardy IR, et al. Switch-mediated activation and retargeting of CAR-T cells for B-cell malignancies. Proc Natl Acad Sci 2016; 113:E459–468. http://doi.org/10.1073/pnas.1524155113.

10. Cao Y, Rodgers DT, Du J, Ahmad I, Hampton EN, Ma JS, et al. Design of Switchable Chimeric Antigen Receptor T Cells Targeting Breast Cancer. Angew Chem Int Ed Engl 2016; 55:7520–7524. http://doi.org/10.1002/anie.201601902.

11. Cartellieri M, Feldmann A, Koristka S, Arndt C, Loff S, Ehninger A, et al. Switching CAR T cells on and off: a novel modular platform for retargeting of T cells to AML blasts. Blood Cancer J 2016; 6:E458. http://doi.org/10.1038/bcj.2016.61.

12. Loureiro LR, Feldmann A, Bergmann R, Koristka S, Berndt N, Arndt C, et al. Development of a novel target module redirecting UniCAR T cells to Sialyl Tn-expressing tumor cells. Blood Cancer J 2018; 8:81. http://doi.org/10.1038/s41408-018-0113-4.

13. Arcangeli S, Rotiroti MC, Bardelli M, Simonelli L, Magnani CF, Biondi A, et al. Balance of Anti-CD123 Chimeric Antigen Receptor Binding Affinity and Density for the Targeting of Acute Myeloid Leukemia. Mol Ther 2017; 25:1933–1945. http://doi.org/10.1016/j.ymthe.2017.04.017.

14. Lynn RC, Weber EW, Sotillo E, Gennert D, Xu P, Good Z, et al. c-Jun overexpression in CAR T cells induces exhaustion resistance. Nature 2019; 576:293–300. http://doi.org/10.1038/s41586-019-1805-z.

15. Lamers CH, Willemsen R, van Elzakker P, van Steenbergen-Langeveld S, Broertjes M, Oosterwijk-Wakka J, et al. Immune responses to transgene and retroviral vector in patients treated with ex vivo-engineered T cells. Blood 2011; 117:72–82. http://doi.org/10.1182/blood-2010-07-294520.

16. Howard SC, Trifilio S, Gregory TK, Baxter N, McBride A. Tumor lysis syndrome in the era of novel and targeted agents in patients with hematologic malignancies: a systematic review. Ann Hematol 2016; 95:563–573. http://doi.org/10.1007/s00277-015-2585-7.

17. Morgan RA, Yang JC, Kitano M, Dudley ME, Laurencot CM, Rosenberg SA. Case report of a serious adverse event following the administration of T cells transduced with a chimeric antigen receptor recognizing ERBB2. Mol Ther 2010; 18:843–851. http://doi.org/10.1038/mt.2010.24.

18. Liu X, Jiang S, Fang C, Yang S, Olalere D, Pequignot EC, et al. Affinity-Tuned ErbB2 or EGFR Chimeric Antigen Receptor T Cells Exhibit an Increased Therapeutic Index against Tumors in Mice. Cancer Res 2015; 75:3596–3607. http://doi.org/10.1158/0008-5472.CAN-15-0159.

19. Drent E, Themeli M, Poels R, de Jong-Korlaar R, Yuan H, de Bruijn J, et al. A Rational Strategy for Reducing On-Target Off-Tumor Effects of CD38-Chimeric Antigen Receptors by Affinity Optimization. Mol Ther 2017; 25:1946–1958. http://doi.org/10.1016/j.ymthe.2017.04.024.

20. Caratelli S, Arriga R, Sconocchia T, Ottaviani A, Lanzilli G, Pastore D, et al. In vitro elimination of epidermal growth factor receptor-overexpressing cancer cells by CD32A-chimeric receptor T cells in combination with cetuximab or panitumumab. Int J Cancer 2020; 146:236–247. http://doi.org/10.1002/ijc.32663.

21. Arriga R, Caratelli S, Lanzilli G, Ottaviani A, Cenciarelli C, Sconocchia T, et al. CD16-158-valine chimeric receptor T cells overcome the resistance of KRAS-mutated colorectal carcinoma cells to cetuximab. Int J Cancer 2020; 146:2531–2538. http://doi.org/10.1002/ijc.32618.

22. Clémenceau B, Vivien R, Pellat C, Foss M, Thibault G, Vié H. The human natural killer cytotoxic cell line NK-92, once armed with a murine CD16 receptor, represents a convenient cellular tool for the screening of mouse mAbs according to their ADCC potential. MAbs 2013; 5:587–594. http://doi.org/10.4161/mabs.25077.

23. Kudo K, Imai C, Lorenzini P, Kamiya T, Kono K, Davidoff AM, et al. T lymphocytes expressing a CD16 signaling receptor exert antibody-dependent cancer cell killing. Cancer Res 2014; 74:93–103. http://doi.org/10.1158/0008-5472.CAN-13-1365.

24. Ochi F, Fujiwara H, Tanimoto K, Asai H, Miyazaki Y, Okamoto S, et al. Gene-modified human α/β-T cells expressing a chimeric CD16-CD3ζ receptor as adoptively transferable effector cells for anticancer monoclonal antibody therapy. Cancer Immunol Res 2014; 2:249–262. http://doi.org/10.1158/2326-6066.CIR-13-0099-T.

25. Clémenceau B, Valsesia-Wittmann S, Jallas AC, Vivien R, Rousseau R, Marabelle A, et al. In Vitro and In Vivo Comparison of Lymphocytes Transduced with a Human CD16 or with a Chimeric Antigen Receptor Reveals Potential Off-Target Interactions due to the IgG2 CH2-CH3 CAR-Spacer. J Immunol Res 2015; 2015:482089. http://doi.org/10.1155/2015/482089.

26. D’Aloia MM, Caratelli S, Palumbo C, Battella S, Arriga R, Lauro D, et al. T lymphocytes engineered to express a CD16-chimeric antigen receptor redirect T-cell immune responses against immunoglobulin G-opsonized target cells. Cytotherapy 2016; 18:278–290. http://doi.org/10.1016/j.jcyt.2015.10.014.

27. Tanaka H, Fujiwara H, Ochi F, Tanimoto K, Casey N, Okamoto S, et al. Development of Engineered T Cells Expressing a Chimeric CD16-CD3ζ Receptor to Improve the Clinical Efficacy of Mogamulizumab Therapy Against Adult T-Cell Leukemia. Clin Cancer Res 2016; 22:4405–4416. http://doi.org/10.1158/1078-0432.CCR-15-2714.

28. Oberg HH, Kellner C, Gonnermann D, Sebens S, Bauerschlag D, Gramatzki M, et al. Tribody [(HER2)2xCD16] Is More Effective Than Trastuzumab in Enhancing γδ T Cell and Natural Killer Cell Cytotoxicity Against HER2-Expressing Cancer Cells. Front Immunol 2018; 9:814. http://doi.org/10.3389/fimmu.2018.00814.

29. Rataj F, Jacobi SJ, Stoiber S, Asang F, Ogonek J, Tokarew N, et al. High-affinity CD16-polymorphism and Fc-engineered antibodies enable activity of CD16-chimeric antigen receptor-modified T cells for cancer therapy. Br J Cancer 2019; 120:79–87. http://doi.org/10.1038/s41416-018-0341-1.

30. Li B, Zhao L, Guo H, Wang C, Zhang X, Wu L, Chen L, Tong Q, Qian W, Wang H, Guo Y. Characterization of a rituximab variant with potent antitumor activity against rituximab-resistant B-cell lymphoma. Blood 2009 Dec 3; 114:5007–15. http://doi.org/10.1182/blood-2009-06-225474.

31. Han L, Chen J, Ding K, Zong H, Xie Y, Jiang H, Zhang B, Lu H, Yin W, Gilly J, Zhu J. Efficient generation of bispecific IgG antibodies by split intein mediated protein trans-splicing system. Sci Rep 2017 Aug 21; 7:8360. http://doi.org/10.1038/s41598-017-08641-3.

32. Long AH, Haso WM, Shern JF, Wanhainen KM, Murgai M, Ingaramo M, et al. 4-1BB costimulation ameliorates T cell exhaustion induced by tonic signaling of chimeric antigen receptors. Nat Med 2015; 21:581–590. http://doi.org/10.1038/nm.3838.

33. Yuan Y, Zong H, Bai J, Han L, Wang L, Zhang X, et al. Bioprocess development of a stable FUT8−/− -CHO cell line to produce defucosylated anti-HER2 antibody. Bioprocess Biosyst Eng 2019; 42:1263–1271. http://doi.org/10.1007/s00449-019-02124-7.

34. Ashoor DN, Ben Khalaf N, Bourguiba-Hachemi S, Marzouq MH, Fathallah MD. Engineering of the upper hinge region of human IgG1 Fc enhances the binding affinity to FcγIIIa (CD16a) receptor isoform. Protein Eng Des Sel 2018; 31:205–212. http://doi.org/10.1093/protein/gzy019.

35. Veeramani S, Wang SY, Dahle C, Blackwell S, Jacobus L, Knutson T, et al. Rituximab infusion induces NK activation in lymphoma patients with the high-affinity CD16 polymorphism. Blood 2011; 118:3347–3349. http://doi.org/10.1182/blood-2011-05-351411.

36. Mazor Y, Van Blarcom T, Mabry R, Iverson BL, Georgiou G. Isolation of engineered, full-length antibodies from libraries expressed in Escherichia coli. Nat Biotechnol 2007; 25:563–565. http://doi.org/10.1038/nbt1296.

37. Dashivets T, Thomann M, Rueger P, Knaupp A, Buchner J, Schlothauer T. Multi-Angle Effector Function Analysis of Human Monoclonal IgG Glycovariants. PLoS One 2015; 10:E0143520. http://doi.org/10.1371/journal.pone.0143520.

38. Hristodorov D, Fischer R, Joerissen H, Müller-Tiemann B, Apeler H, Linden L. Generation and comparative characterization of glycosylated and aglycosylated human IgG1 antibodies. Mol Biotechnol 2013; 53:326–335. http://doi.org/10.1007/s12033-012-9531-x.

39. Sheth VS, Gauthier J. Taming the beast: CRS and ICANS after CAR T-cell therapy for ALL. Bone Marrow Transplant 2021; 56:552–566. http://doi.org/10.1038/s41409-020-01134-4.

40. Weber R, Groth C, Lasser S, Arkhypov I, Petrova V, Altevogt P, et al. IL-6 as a major regulator of MDSC activity and possible target for cancer immunotherapy. Cell Immunol 2021; 359:104254. http://doi.org/10.1016/j.cellimm.2020.104254.

41. Mittal SK, Cho KJ, Ishido S, Roche PA. Interleukin 10 (IL-10)-mediated Immunosuppression: MARCH-I INDUCTION REGULATES ANTIGEN PRESENTATION BY MACROPHAGES BUT NOT DENDRITIC CELLS. J Biol Chem 2015; 290:27158–27167. http://doi.org/10.1074/jbc.M115.682708.

42. Ghorashian S, Kramer AM, Onuoha S, Wright G, Bartram J, Richardson R, et al. Enhanced CAR T cell expansion and prolonged persistence in pediatric patients with ALL treated with a low-affinity CD19 CAR. Nat Med 2019; 25:1408–1414. http://doi.org/10.1038/s41591-019-0549-5.

43. Rudnick SI, Lou J, Shaller CC, Tang Y, Klein-Szanto AJ, Weiner LM, Marks JD, Adams GP. Influence of affinity and antigen internalization on the uptake and penetration of Anti-HER2 antibodies in solid tumors. Cancer Res 2011 Mar 15; 71:2250–9. http://doi.org/10.1158/0008-5472.CAN-10-2277.

44. Thurber GM, Schmidt MM, Wittrup KD. Factors determining antibody distribution in tumors. Trends Pharmacol Sci 2008 Feb; 29:57–61. http://doi.org/10.1016/j.tips.2007.11.004.

45. van Mirre E, Teeling JL, van der Meer JW, Bleeker WK, Hack CE. Monomeric IgG in intravenous Ig preparations is a functional antagonist of FcgammaRII and FcgammaRIIIb. J Immunol 2004; 173:332–339. http://doi.org/10.4049/jimmunol.173.1.332‥

46. Lux A, Yu X, Scanlan CN, Nimmerjahn F. Impact of immune complex size and glycosylation on IgG binding to human FcγRs. J Immunol 2013; 190:4315–4323. http://doi.org/10.4049/jimmunol.1200501.

47. Koenderman L. Inside-Out Control of Fc-Receptors. Front Immunol 2019; 10:544. http://doi.org.10.3389/fimmu.2019.00544.

48. Brandsma AM, Schwartz SL, Wester MJ, Valley CC, Blezer GLA, Vidarsson G, et al. Mechanisms of inside-out signaling of the high-affinity IgG receptor FcγRI. Sci Signal 2018; 11:eaaq0891. http://doi:.org/10.1126/scisignal.aaq0891.

