## Supplemental files for "Universal chimeric Fcγ receptor T cells with appropriate affinity for IgG1 antibody exhibit optimal antitumor efficacy"

**Supplementary data**

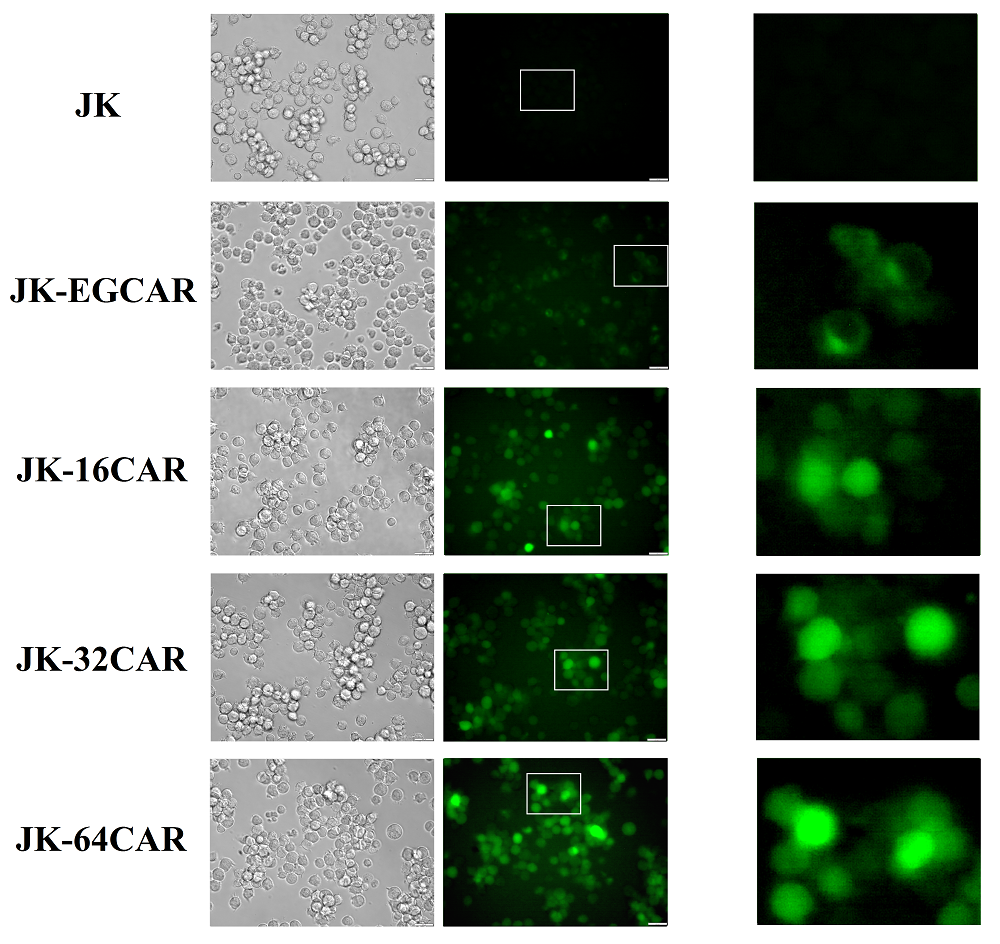

**Figure S1**. The detection of GFP expression with a fluorescence microscope. The cells were analyzed 80 h after lentivirus infection.

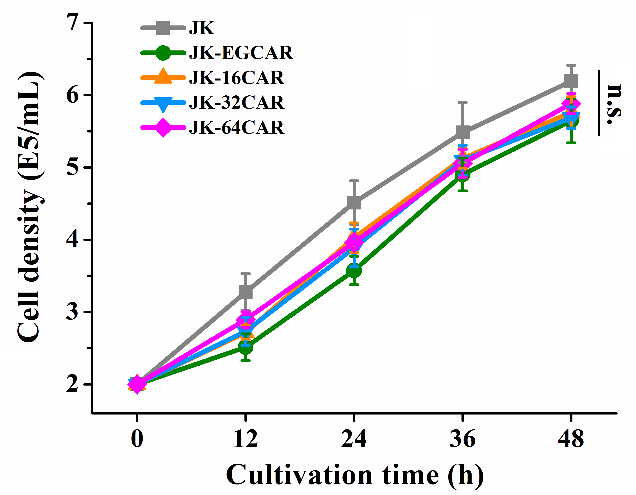

**Figure S2**. Cell proliferation activities of the Jurkat-EG/16/32/64CAR and primary Jurkat cells. The cell densities were set as 2E5/mL and measured every 12 h. Data are presented as mean ± SD, n = 3. Statistical analysis was based on two-tailed heteroscedastic Student’s t-test, and statistical significance is considered at p<0.05. “n.s.” indicates not significant (p>0.05).

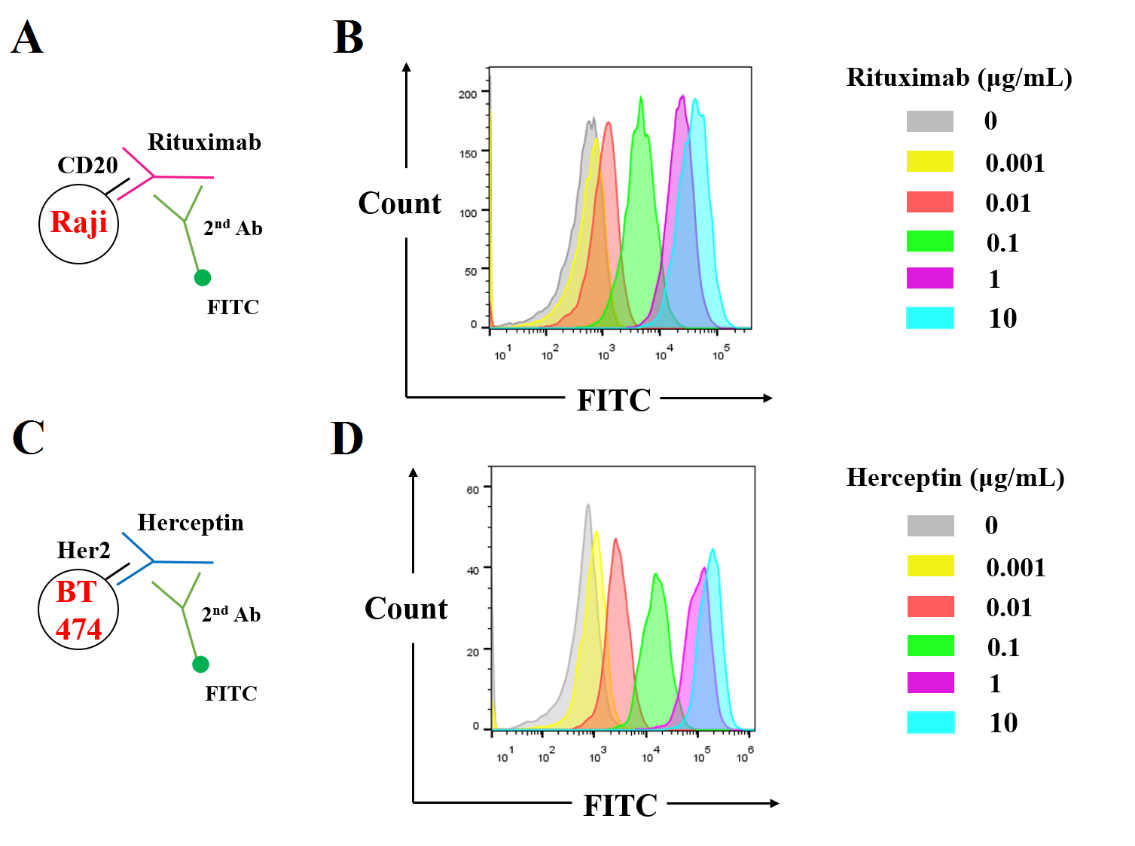

**Figure S3**. The expression levels of the two antigens on the target cells. FACS analysis of CD20 on Raji cells (**A**, **B**) and Her2 on BT474 cells (**C**, **D**). Raji cells (1E6/mL)/BT474 cells (3E5/mL) were incubated with different concentrations of rituximab/herceptin and goat anti-human IgG antibody (FITC) in sequence, and then cell staining was measured by FACS.

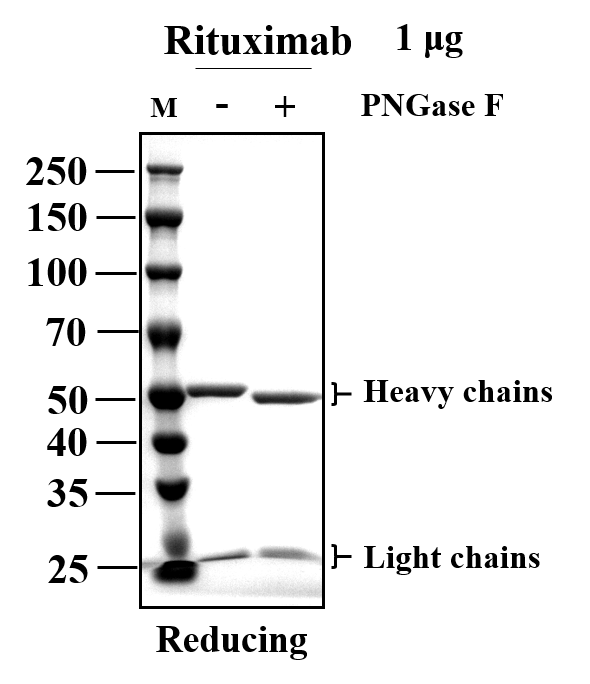

**Figure S4**. Verification of the de-glycolysation by SDS-PAGE. Rituximab was digested by PNGase F at 37 °C for 24 h under native conditions. The digestion products were sampled and analyzed by reducing SDS-PAGE.

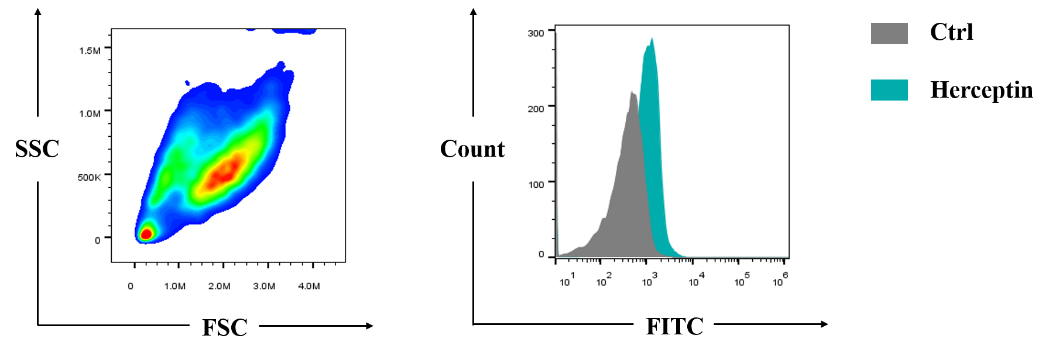

**Figure S5**. FACS analysis of the expression of Her2 on the transplanted U251 MG tumor cells. Female nude mice were implanted s.c. with 5E6 U251 MG cells on day0, and the tumor was collected on day8. The single-cell suspensions of tumor tissue were prepared and incubated with herceptin and goat anti-human IgG antibody (FITC) in sequence. After washing twice with PBS, the cells were analyzed by FACS.

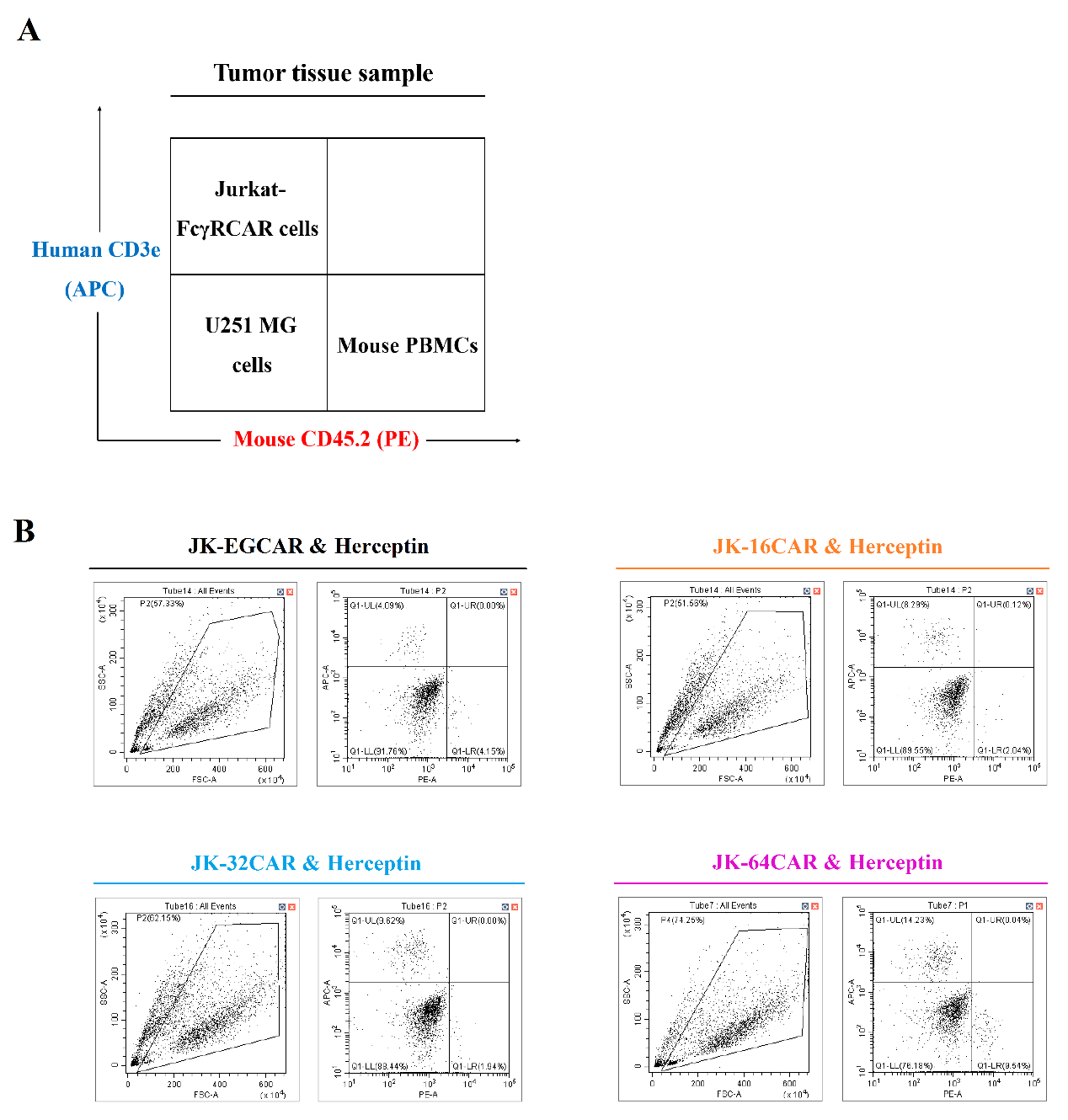

**Figure S6**. FACS analysis of the tumor tissue samples (U251 MG model). (**A**) Illustration of the cell distribution. (**B**) Results of the FACS analysis. The single-cell suspensions of tumor tissues were incubated with Mouse TruStain FcX and Human TruStain FcX, subsequently anti-mouse CD45.2 antibody (PE) and anti-human CD3e antibody (APC). After washing twice with PBS, the cells were analyzed by FACS.

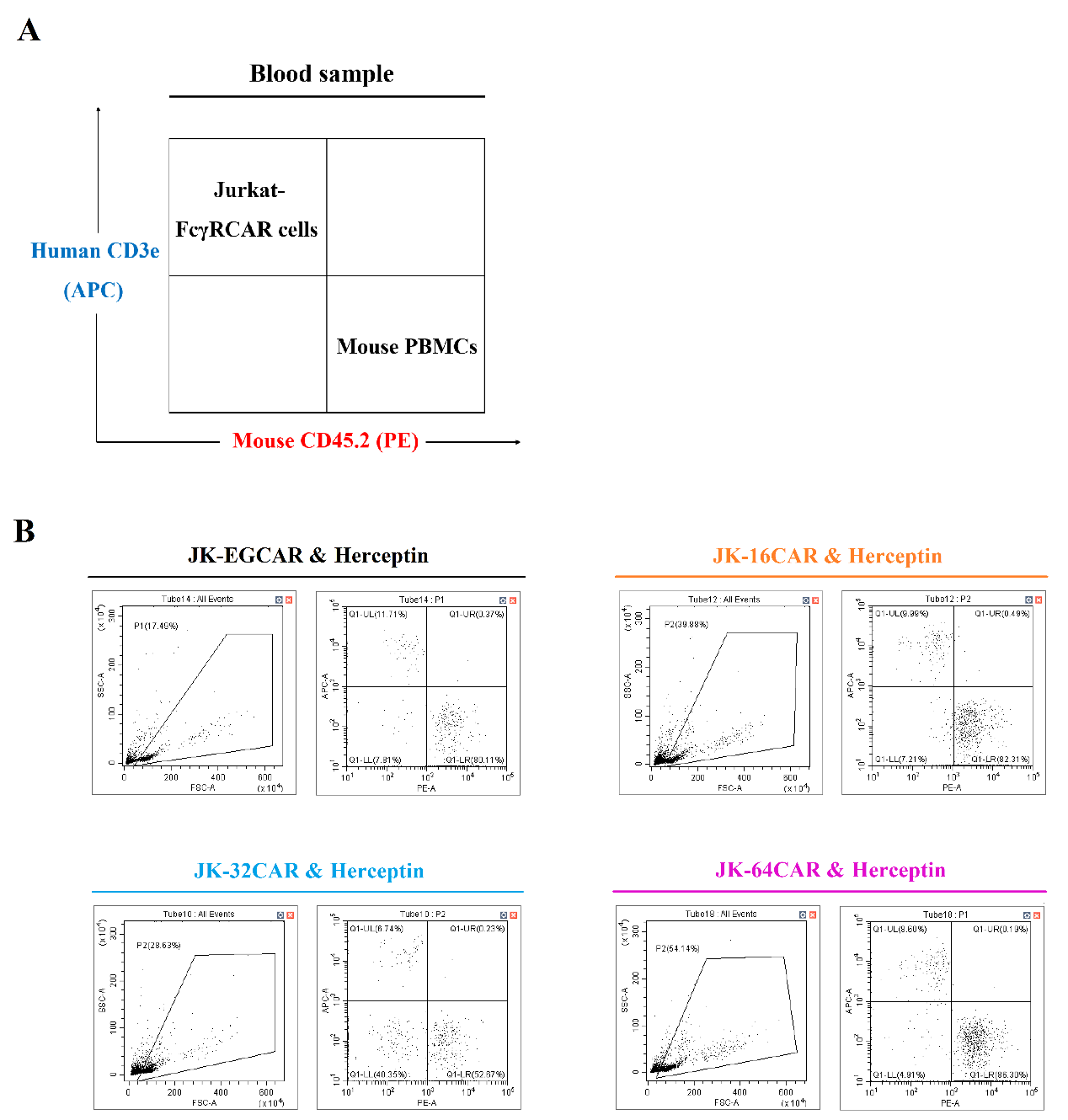

**Figure S7**. FACS analysis of the blood samples (U251 MG model). (**A**) Illustration of the cell distribution. (**B**) Results of the FACS analysis. The separated PBMCs were incubated with Mouse TruStain FcX and Human TruStain FcX, subsequently anti-mouse CD45.2 antibody (PE) and anti-human CD3e antibody (APC). After washing twice with PBS, the cells were analyzed by FACS.

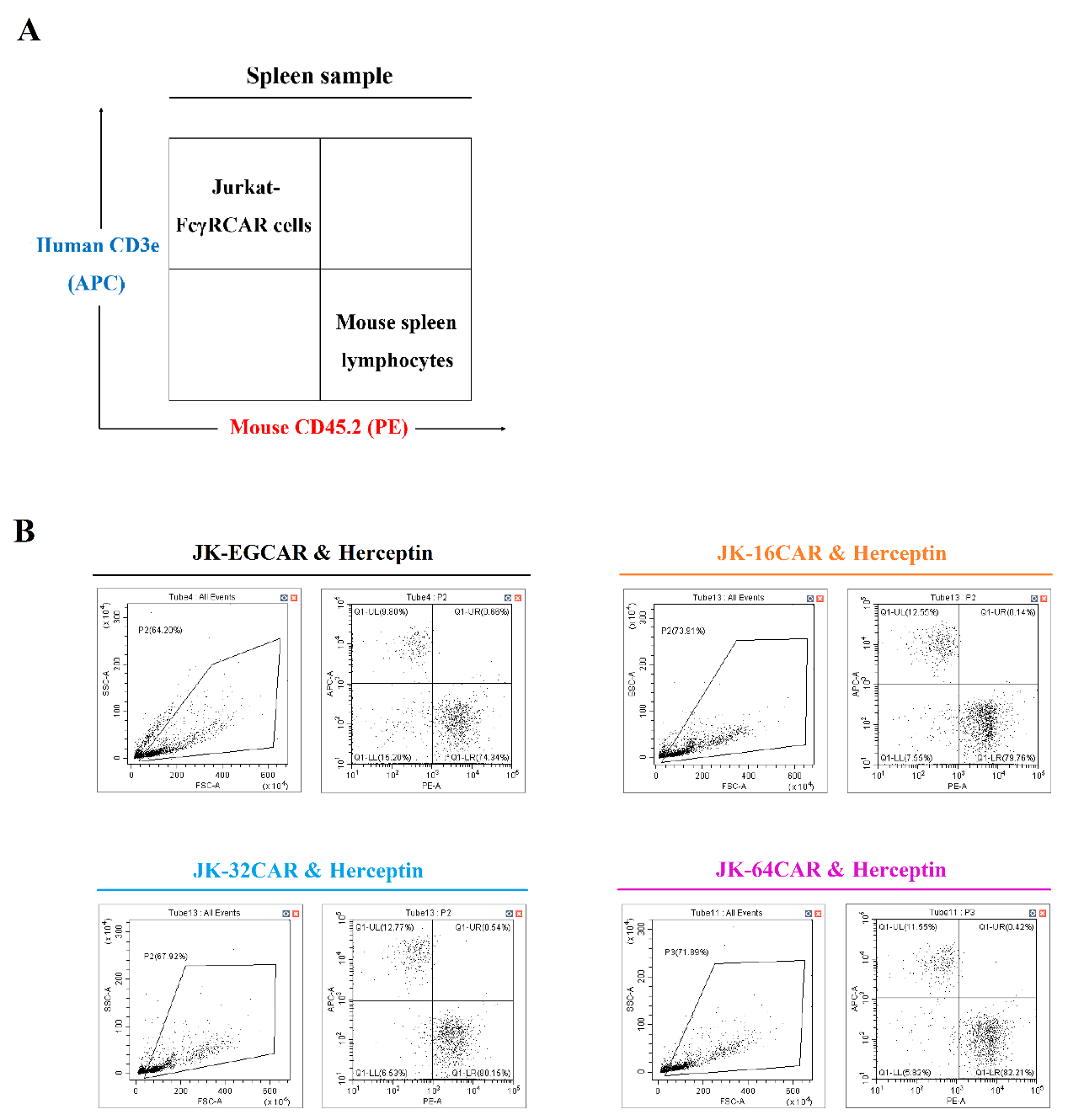

**Figure S8**. FACS analysis of the spleen samples (U251 MG model). (**A**) Illustration of the cell distribution. (**B**) Results of the FACS analysis. The separated spleen lymphocytes were incubated with Mouse TruStain FcX and Human TruStain FcX, subsequently anti-mouse CD45.2 antibody (PE) and anti-human CD3e antibody (APC). After washing twice with PBS, the cells were analyzed by FACS.

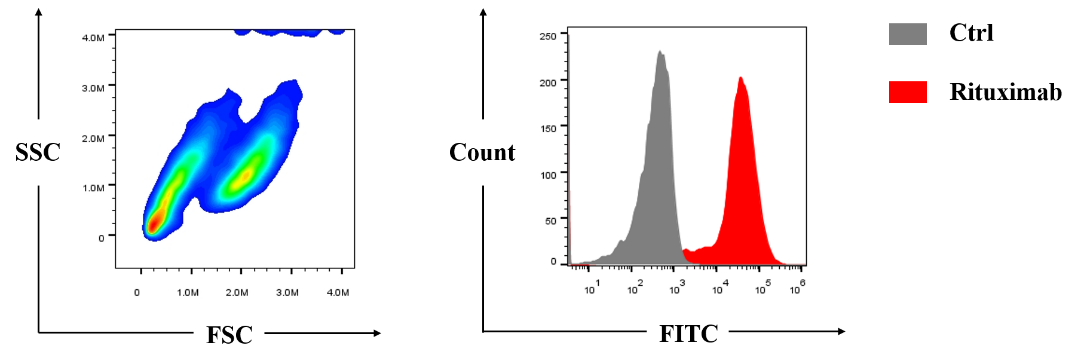

**Figure S9**. FACS analysis of the expression of CD20 on the transplanted Raji tumor cells. Female nude mice were implanted s.c. with 4E6 Raji cells on day0, and the tumor was collected on day10. The single-cell suspensions of tumor tissue were prepared and incubated with rituximab and goat anti-human IgG antibody (FITC) in sequence. After washing twice with PBS, the cells were analyzed by FACS.

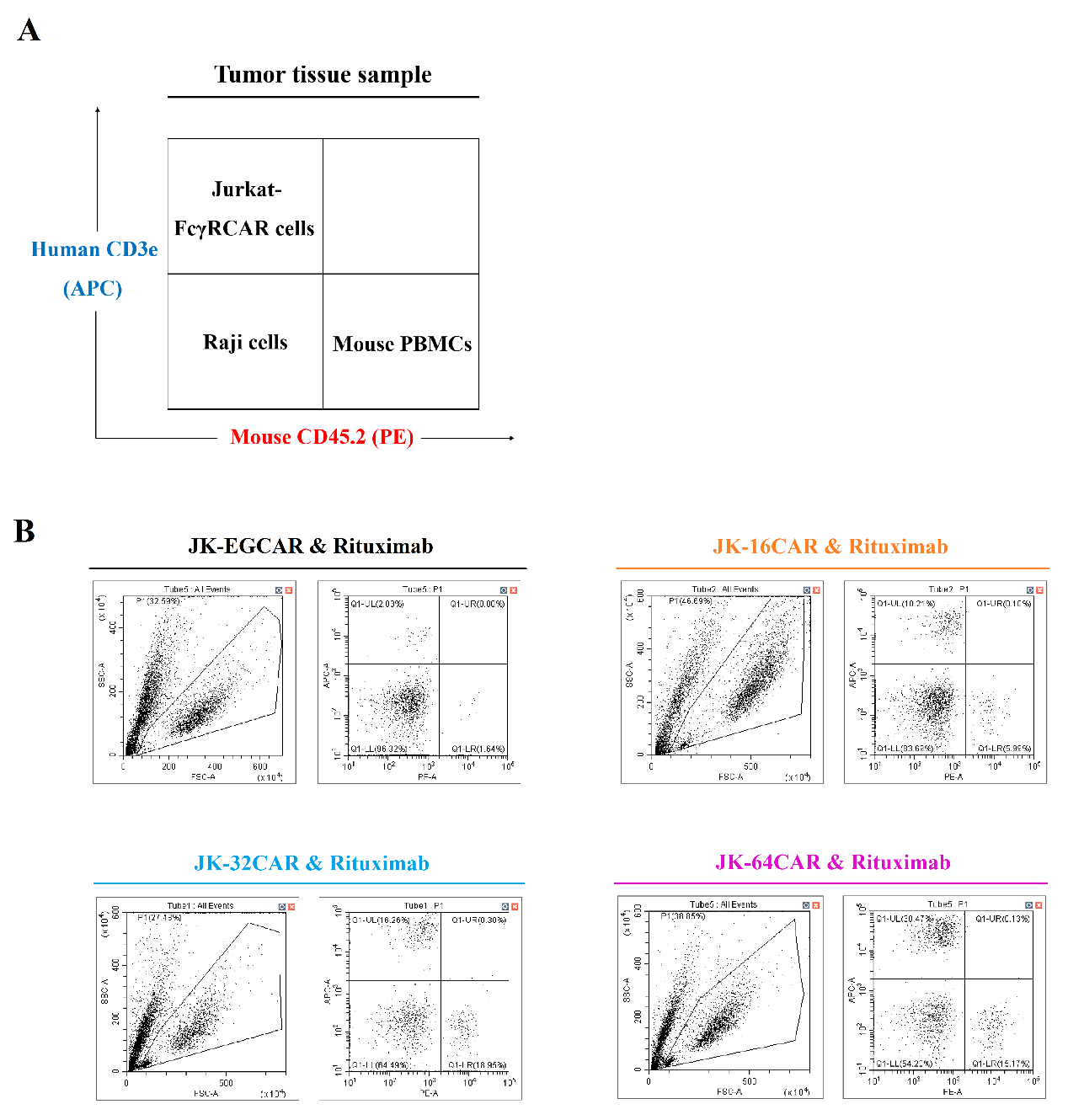

**Figure S10**. FACS analysis of the tumor tissue samples (Raji model). (**A**) Illustration of the cell distribution. (**B**) Results of the FACS analysis. The single-cell suspensions of tumor tissues were incubated with Mouse TruStain FcX and Human TruStain FcX, subsequently anti-mouse CD45.2 antibody (PE) and anti-human CD3e antibody (APC). After washing twice with PBS, the cells were analyzed by FACS.

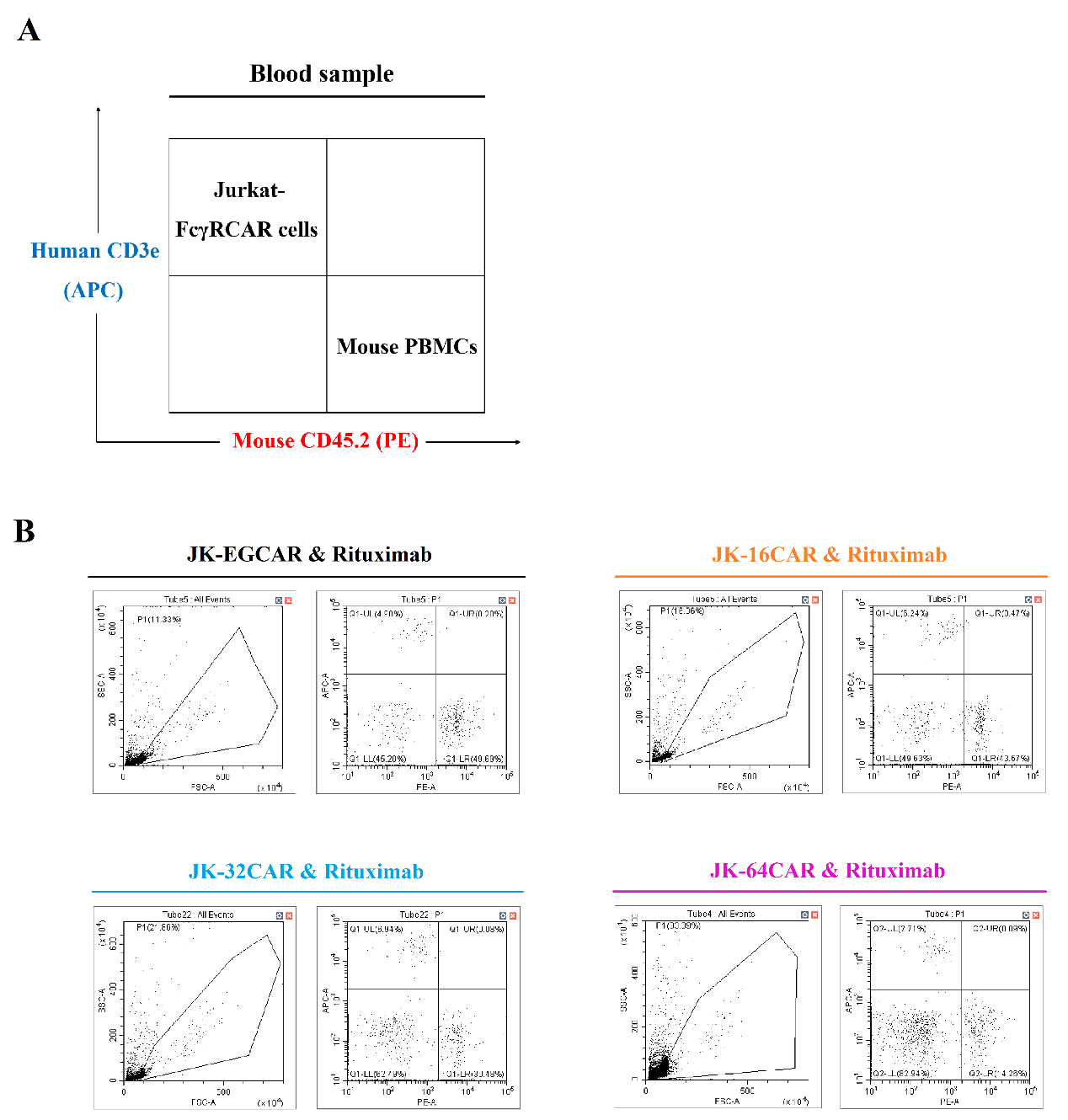

**Figure S11**. FACS analysis of the blood samples (Raji model). (**A**) Illustration of the cell distribution. (**B**) Results of the FACS analysis. The separated PBMCs were incubated with Mouse TruStain FcX and Human TruStain FcX, subsequently anti-mouse CD45.2 antibody (PE) and anti-human CD3e antibody (APC). After washing twice with PBS, the cells were analyzed by FACS.

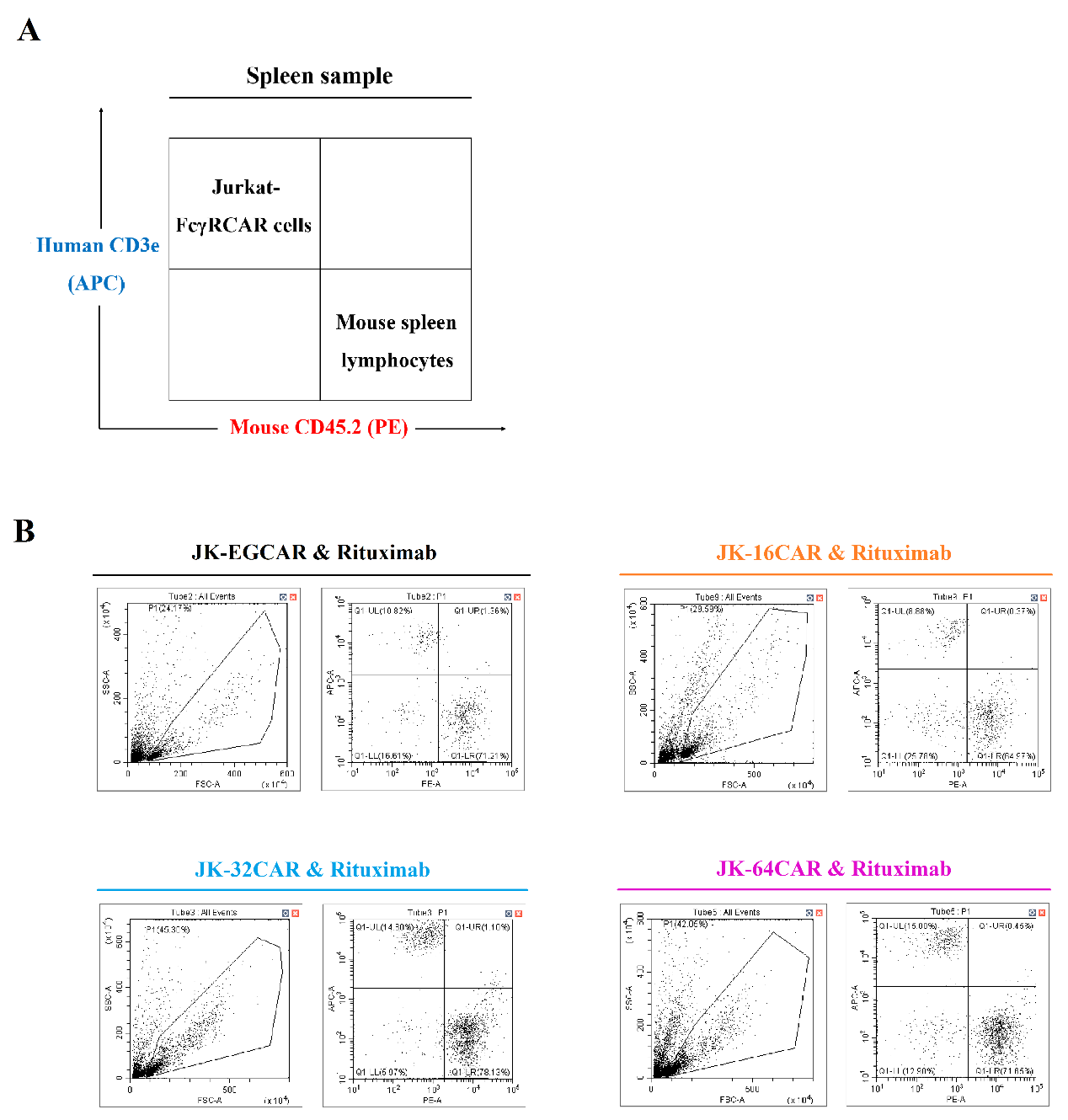

**Figure S12**. FACS analysis of the spleen samples (Raji model). (**A**) Illustration of the cell distribution. (**B**) Results of the FACS analysis. The separated spleen lymphocytes were incubated with Mouse TruStain FcX and Human TruStain FcX, subsequently anti-mouse CD45.2 antibody (PE) and anti-human CD3e antibody (APC). After washing twice with PBS, the cells were analyzed by FACS.

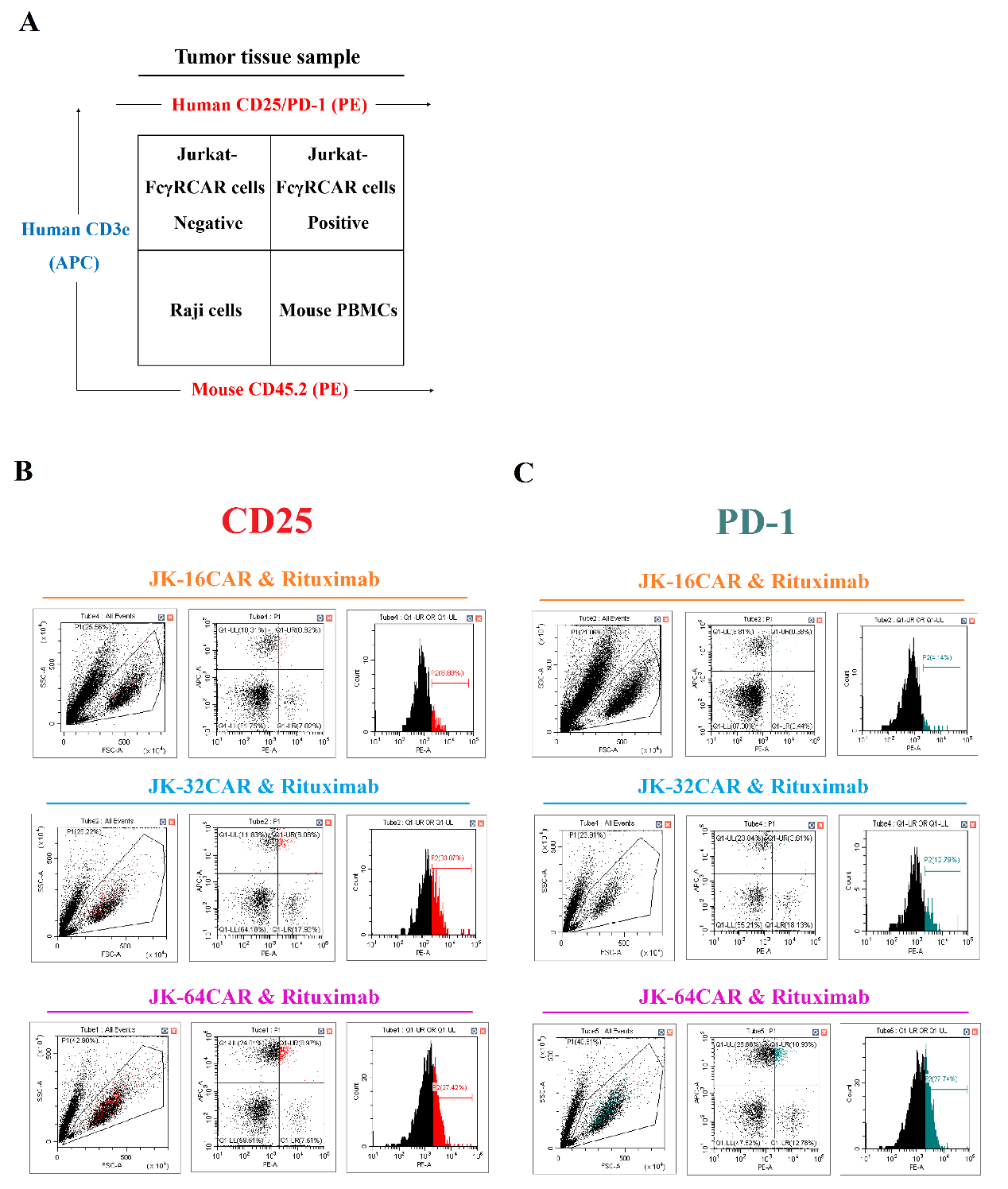

**Figure S13**. FACS analysis of the immune status (Raji model). (**A**) Illustration of the cell distribution. (**B**) Results of the FACS analysis. The single-cell suspensions of tumor tissues were incubated with Mouse TruStain FcX and Human TruStain FcX, subsequently anti-mouse CD45.2 antibody (PE), anti-human CD3e antibody (APC), and anti-human CD25 antibody (PE) or anti-human PD-1 antibody (PE). After washing twice with PBS, the cells were analyzed by FACS.

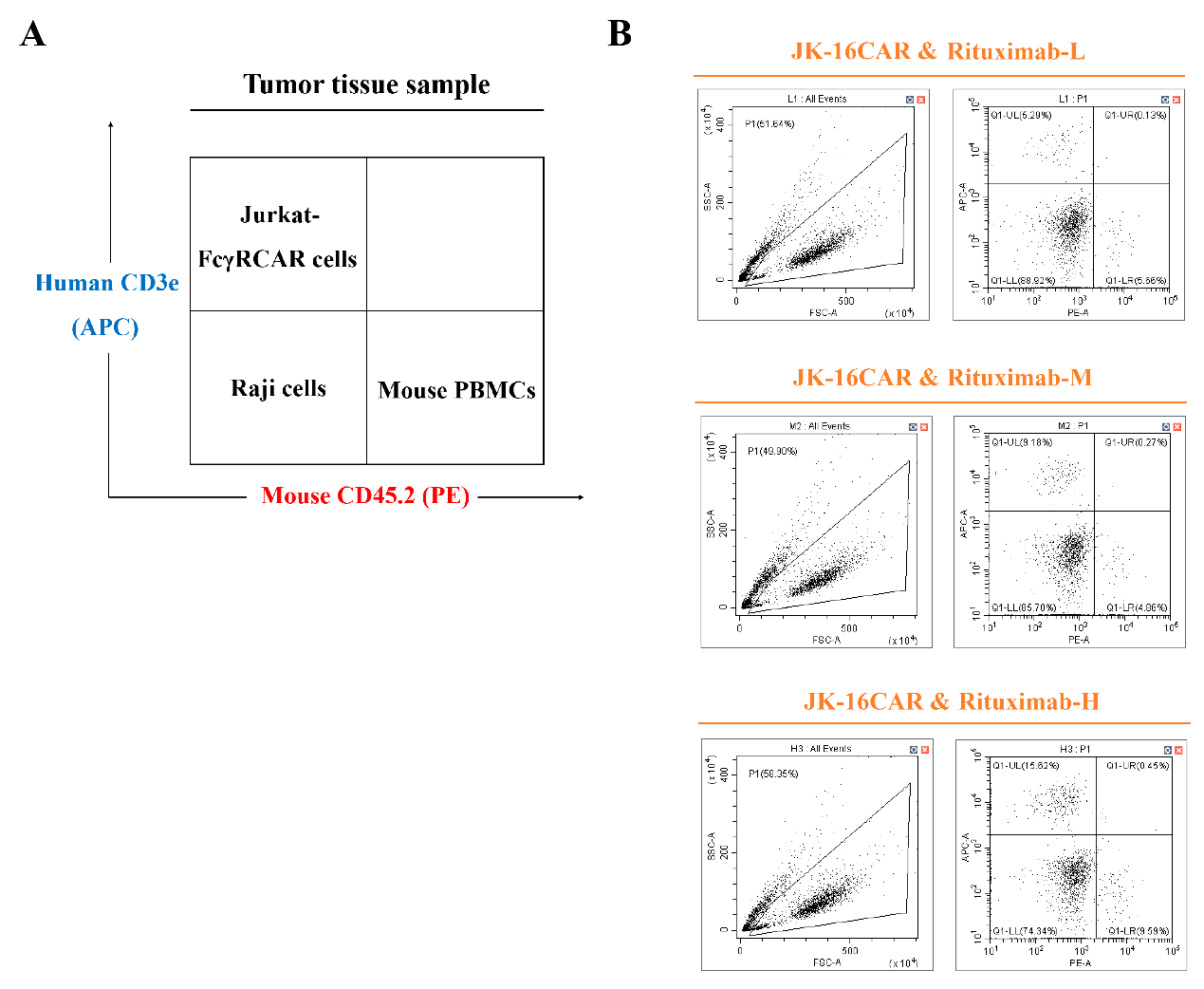

**Figure S14**. FACS analysis of the tumor tissue samples (Rituximab mutants model). (**A**) Illustration of the cell distribution. (**B**) Results of the FACS analysis. The single-cell suspensions of tumor tissues were incubated with Mouse TruStain FcX and Human TruStain FcX, subsequently anti-mouse CD45.2 antibody (PE) and anti-human CD3e antibody (APC). After washing twice with PBS, the cells were analyzed by FACS.

Gene sequence of CD8α signal peptide:

atggccttaccagtgaccgccttgctcctgccgctggccttgctgctccacgccgccaggccg

Amino acid sequence of CD8α signal peptide:

MALPVTALLLPLALLLHAARP

Gene sequence of EGFP:

atggtgagcaagggcgaggagctgttcaccggggtggtgcccatcctggtcgagctggacggcgacgtaaacggccacaagttcagcgtgtccggcgagggcgagggcgatgccacctacggcaagctgaccctgaagttcatctgcaccaccggcaagctgcccgtgccctggcccaccctcgtgaccaccctgacctacggcgtgcagtgcttcagccgctaccccgaccacatgaagcagcacgacttcttcaagtccgccatgcccgaaggctacgtccaggagcgcaccatcttcttcaaggacgacggcaactacaagacccgcgccgaggtgaagttcgagggcgacaccctggtgaaccgcatcgagctgaagggcatcgacttcaaggaggacggcaacatcctggggcacaagctggagtacaactacaacagccacaacgtctatatcatggccgacaagcagaagaacggcatcaaggtgaacttcaagatccgccacaacatcgaggacggcagcgtgcagctcgccgaccactaccagcagaacacccccatcggcgacggccccgtgctgctgcccgacaaccactacctgagcacccagtccgccctgagcaaagaccccaacgagaagcgcgatcacatggtcctgctggagttcgtgaccgccgccgggatcactctcggcatggacgagctgtacaag

Amino acid sequence of EGFP:

MVSKGEELFTGVVPILVELDGDVNGHKFSVSGEGEGDATYGKLTLKFICTTGKLPVPWPTLVTTLTYGVQCFSRYPDHMKQHDFFKSAMPEGYVQERTIFFKDDGNYKTRAEVKFEGDTLVNRIELKGIDFKEDGNILGHKLEYNYNSHNVYIMADKQKNGIKVNFKIRHNIEDGSVQLADHYQQNTPIGDGPVLLPDNHYLSTQSALSKDPNEKRDHMVLLEFVTAAGITLGMDELYK

Gene sequence of CD16a outer membrane domain:

ggcatgcggactgaagatctcccaaaggctgtggtgttcctggagcctcaatggtacagggtgctcgagaaggacagtgtgactctgaagtgccagggagcctactcccctgaggacaattccacacagtggtttcacaatgagagcctcatctcaagccaggcctcgagctacttcattgacgctgccacagtcgacgacagtggagagtacaggtgccagacaaacctctccaccctcagtgacccggtgcagctagaagtccatatcggctggctgttgctccaggcccctcggtgggtgttcaaggaggaagaccctattcacctgaggtgtcacagctggaagaacactgctctgcataaggtcacatatttacagaatggcaaaggcaggaagtattttcatcataattctgacttctacattccaaaagccacactcaaagacagcggctcctacttctgcagggggctttttgggagtaaaaatgtgtcttcagagactgtgaacatcaccatcactcaaggtttggcagtgtcaaccatctcatcattctttccacctgggtaccaa

Amino acid sequence of CD16a outer membrane domain:

GMRTEDLPKAVVFLEPQWYRVLEKDSVTLKCQGAYSPEDNSTQWFHNESLISSQASSYFIDAATVDDSGEYRCQTNLSTLSDPVQLEVHIGWLLLQAPRWVFKEEDPIHLRCHSWKNTALHKVTYLQNGKGRKYFHHNSDFYIPKATLKDSGSYFCRGLFGSKNVSSETVNITITQGLAVSTISSFFPPGYQ

Gene sequence of CD32a outer membrane domain:

caagctgcagctcccccaaaggctgtgctgaaacttgagcccccgtggatcaacgtgctccaggaggactctgtgactctgacatgccagggggctcgcagccctgagagcgactccattcagtggttccacaatgggaatctcattcccacccacacgcagcccagctacaggttcaaggccaacaacaatgacagcggggagtacacgtgccagactggccagaccagcctcagcgaccctgtgcatctgactgtgctttccgaatggctggtgctccagacccctcacctggagttccaggagggagaaaccatcatgctgaggtgccacagctggaaggacaagcctctggtcaaggtcacattcttccagaatggaaaatcccagaaattctcccgtttggatcccaccttctccatcccacaagcaaaccacagtcacagtggtgattaccactgcacaggaaacataggctacacgctgttctcatccaagcctgtgaccatcactgtccaagtgcccagcatgggcagctcttcaccaatgggg

Amino acid sequence of CD32a outer membrane domain:

QAAAPPKAVLKLEPPWINVLQEDSVTLTCQGARSPESDSIQWFHNGNLIPTHTQPSYRFKANNNDSGEYTCQTGQTSLSDPVHLTVLSEWLVLQTPHLEFQEGETIMLRCHSWKDKPLVKVTFFQNGKSQKFSRLDPTFSIPQANHSHSGDYHCTGNIGYTLFSSKPVTITVQVPSMGSSSPMG

Gene sequence of CD64 outer membrane domain:

caagtggacaccacaaaggcagtgatcactttgcagcctccatgggtcagcgtgttccaagaggaaaccgtaaccttgcactgtgaggtgctccatctgcctgggagcagctctacacagtggtttctcaatggcacagccactcagacctcgacccccagctacagaatcacctctgccagtgtcaatgacagtggtgaatacaggtgccagagaggtctctcagggcgaagtgaccccatacagctggaaatccacagaggctggctactactgcaggtctccagcagagtcttcacggaaggagaacctctggccttgaggtgtcatgcgtggaaggataagctggtgtacaatgtgctttactatcgaaatggcaaagcctttaagtttttccactggaattctaacctcaccattctgaaaaccaacataagtcacaatggcacctaccattgctcaggcatgggaaagcatcgctacacatcagcaggaatatctgtcactgtgaaagagctatttccagctccagtgctgaatgcatctgtgacatccccactcctggaggggaatctggtcaccctgagctgtgaaacaaagttgctcttgcagaggcctggtttgcagctttacttctccttctacatgggcagcaagaccctgcgaggcaggaacacatcctctgaataccaaatactaactgctagaagagaagactctgggttatactggtgcgaggctgccacagaggatggaaatgtccttaagcgcagccctgagttggagcttcaagtgcttggcctccagttaccaactcctgtctggtttcat

Amino acid sequence of CD64 outer membrane domain:

QVDTTKAVITLQPPWVSVFQEETVTLHCEVLHLPGSSSTQWFLNGTATQTSTPSYRITSASVNDSGEYRCQRGLSGRSDPIQLEIHRGWLLLQVSSRVFTEGEPLALRCHAWKDKLVYNVLYYRNGKAFKFFHWNSNLTILKTNISHNGTYHCSGMGKHRYTSAGISVTVKELFPAPVLNASVTSPLLEGNLVTLSCETKLLLQRPGLQLYFSFYMGSKTLRGRNTSSEYQILTARREDSGLYWCEAATEDGNVLKRSPELELQVLGLQLPTPVWFH

Gene sequence of CD28 hinge domain:

attgaagttatgtatcctcctccttacctagacaatgagaagagcaatggaaccattatccatgtgaaagggaaacacctttgtccaagtcccctatttcccggaccttctaagccc

Amino acid sequence of CD28 hinge domain:

IEVMYPPPYLDNEKSNGTIIHVKGKHLCPSPLFPGPSKP

Gene sequence of CD28 transmembrane domain:

ttttgggtgctggtggtggttggtggagtcctggcttgctatagcttgctagtaacagtggcctttattattttctgggtg

Amino acid sequence of CD28 transmembrane domain:

FWVLVVVGGVLACYSLLVTVAFIIFWV

Gene sequence of CD28 costimulatory domain:

aggagtaagaggagcaggctcctgcacagtgactacatgaacatgactccccgccgccccgggcccacccgcaagcattaccagccctatgccccaccacgcgacttcgcagcctatcgctcc

Amino acid sequence of CD28 costimulatory domain:

RSKRSRLLHSDYMNMTPRRPGPTRKHYQPYAPPRDFAAYRS

Gene sequence of CD137 costimulatory domain:

aaacggggcagaaagaaactcctgtatatattcaaacaaccatttatgagaccagtacaaactactcaagaggaagatggctgtagctgccgatttccagaagaagaagaaggaggatgtgaactg

Amino acid sequence of CD137 costimulatory domain:

KRGRKKLLYIFKQPFMRPVQTTQEEDGCSCRFPEEEEGGCEL

Gene sequence of CD247 costimulatory domain:

agagtgaagttcagcaggagcgcagacgcccccgcgtaccagcagggccagaaccagctctataacgagctcaatctaggacgaagagaggagtacgatgttttggacaagagacgtggccgggaccctgagatggggggaaagccgcagagaaggaagaaccctcaggaaggcctgtacaatgaactgcagaaagataagatggcggaggcctacagtgagattgggatgaaaggcgagcgccggaggggcaaggggcacgatggcctttaccagggtctcagtacagccaccaaggacacctacgacgcccttcacatgcaggccctgccccctcgc

Amino acid sequence of CD247 costimulatory domain:

RVKFSRSADAPAYQQGQNQLYNELNLGRREEYDVLDKRRGRDPEMGGKPQRRKNPQEGLYNELQKDKMAEAYSEIGMKGERRRGKGHDGLYQGLSTATKDTYDALHMQALPPR

Gene sequence of P2A peptide:

ggaagcggagccacgaacttctctctgttaaagcaagcaggagatgttgaagaaaaccccgggcct

Amino acid sequence of P2A peptide:

GSGATNFSLLKQAGDVEENPGP
